# Hepatocyte FGF21 is not required for fasting-induced metabolic responses but guides protein appetite after energy depletion

**DOI:** 10.1101/2025.10.22.683899

**Authors:** Justine Bruse, Clothilde Marbach, Arnaud Polizzi, Tatiana Landre, Juliette Salvi, Valérie Alquier-Bacquie, Shiou-Ping Chen, Marine Huillet, Clémence Rives, Céline M.P. Martin, Prunelle Perrier, Fadila Benhamed, Marion Régnier, Stefan Weger, Claire Naylies, Yannick Lippi, Caroline Sommer, Mikael Albin, Frédéric Lasserre, Thierry Levade, Michael Schupp, Laurence Gamet-Payrastre, Léon Kautz, Nicolas Loiseau, Walter Wahli, Sandrine Ellero-Simatos, Céline Cruciani-Guglielmacci, Alexandra Montagner, Alexandre Benani, Catherine Postic, Hervé Guillou, Anne Fougerat

## Abstract

Fasting initiates a coordinated metabolic response to preserve energy balance. As glycogen stores are depleted, the body transitions to mobilizing fatty acids from adipose tissue and generating ketone bodies in the liver to sustain the function of vital organs. A network of hormonal signals and transcriptional programs coordinate these adaptations. Among these, the hepatokine fibroblast growth factor 21 (FGF21) is strongly upregulated during fasting and has been proposed as a key mediator of the fasting response. To investigate the physiological functions of FGF21, we studied mice with hepatocyte-specific deletion of *Fgf21*. Although the liver is the primary source of circulating FGF21 during fasting, its absence in hepatocytes did not alter typical fasting-induced gene expression or key metabolic pathways such as hepatic gluconeogenesis, adipose tissue lipolysis, or ketone production. Instead, we uncovered a distinct role for FGF21 in promoting protein appetite following a fast. These findings challenge the conventional view of hepatocyte-produced FGF21 as a fasting-acting hormone and reveal a more specialized function in guiding nutrient selection after energy depletion.

## Introduction

Adaptation to fluctuations in nutrient availability is fundamental to the survival of the organism and its metabolic health. In mammals, fasting triggers a complex physiological response that involves coordinated transcriptional and metabolic reprogramming across multiple tissues, including the liver (Fougerat *et al*., 2024) and white adipose tissue (WAT) (Ruppert and Kersten, 2025). These adaptations serve to maintain energy homeostasis by shifting metabolic priorities from glucose utilization toward lipid mobilization and ketogenesis (Puchalska and Crawford, 2017). They are driven by hormonal changes and nutrient-sensing transcriptional regulators (Goldstein and Hager, 2015) such as peroxisome proliferator-activated receptor alpha (PPARα) (Kersten *et al*., 1999; Montagner *et al*., 2016), cAMP response element-binding protein (CREB) (Herzig *et al*., 2001; Altarejos and Montminy, 2011), glucocorticoid receptor (GR), and forkhead box protein O1 (FOXO1) (Zhang *et al*., 2006; Haeusler *et al*., 2014). The liver plays a central role in these processes, acting both as a hub and as a source of metabolic and endocrine signals that influence the function of peripheral tissues (Fougerat *et al*., 2024).

One of the most prominently induced hepatic genes during fasting is fibroblast growth factor 21 (*Fgf21*), encoding for the liver-derived hormone or hepatokine FGF21 (Badman *et al*., 2007; Inagaki *et al*., 2007; Gälman *et al*., 2008). Hepatocyte FGF21 expression is strongly upregulated in response to fasting in a PPARα-dependent manner (Badman *et al*., 2007; Inagaki *et al*., 2007; Lundåsen *et al*., 2007; Montagner *et al*., 2016). This rise in FGF21 expression and plasma levels originates from hepatocytes (Markan *et al*., 2014; Montagner *et al*., 2016) and occurs in response to fasting-induced adipose tissue lipolysis (Jaeger *et al*., 2015; Fougerat *et al*., 2022). Early studies suggested that FGF21 functions as a key effector of the metabolic fasting response, promoting ketogenesis, increasing fatty acid oxidation (Badman *et al*., 2007, 2009; Inagaki *et al*., 2007; Potthoff *et al*., 2009) and maintaining glucose homeostasis (Liang *et al*., 2014). Despite these proposed roles, the specific physiological significance of hepatocyte FGF21 during fasting remains incompletely understood because most of these studies were performed using whole body FGF21 knockout mice. In this context, it is worth mentioning that FGF21 is also expressed in many tissues such as the WAT (Dutchak *et al*., 2012), the brown adipose tissue (BAT) (Fisher *et al*., 2012) and the exocrine pancreas (Coate *et al*., 2017). Moreover, while it was initially considered to be a starvation hormone, liver FGF21 has now been reported to be more broadly regulated by many forms of metabolic stresses such as high carbohydrates or alcohol consumption (Talukdar *et al*., 2016; Iroz *et al*., 2017; Song *et al*., 2018; Jensen-Cody *et al*., 2020), ketogenic diet (Badman *et al*., 2007; Song *et al*., 2018), and protein restriction (Laeger *et al*., 2014; Solon-Biet *et al*., 2023).

Although pharmacological or transgenic overexpression of FGF21 improves metabolic outcomes in models of obesity and insulin resistance (Kliewer and Mangelsdorf, 2019), these effects may not accurately reflect the endogenous role of FGF21 under physiological fasting conditions. In the present study, we conducted a comprehensive analysis of the genomic and metabolic responses to fasting in wild-type mice and mice with hepatocyte-restricted deletion of *Fgf21*. Changes of gene expression and metabolic profiling in the liver, WATs, and BAT were assessed. Surprisingly, while fasting induced significant changes in gene expression, we found that the absence of hepatocyte FGF21 had minimal impact on fasting-induced remodeling across these tissues. Instead, we show that hepatocyte FGF21 is required for protein preference following fasting. Our findings challenge the classical concept of hepatocyte-produced FGF21 as a fasting-acting hormone and reveal its role as a key endocrine signal guiding nutrient selection after food deprivation.

## Results and Discussion

### 1. Hepatocyte FGF21 is dispensable for fasting-induced hypoglycemia and ketone body production

Several studies have suggested that liver-derived FGF21 plays a crucial role in the metabolic responses to fasting (Badman *et al*., 2007; Inagaki *et al*., 2007; Potthoff *et al*., 2009; Liang *et al*., 2014). However, most of these reports were based on observations in global *Fgf21* knockout or FGF21 overexpression models, which may not reflect the specific contribution of endogenous hepatic FGF21.

To investigate the physiological function of hepatocyte-produced FGF21, we generated mice with hepatocyte-specific deletion of *Fgf21* (*Fgf21^flox/flox^albumin-Cre^+/−^*, designated as *Fgf21^hep-/-^*) by crossing C57Bl/6J *Fgf21^flox/flox^* mice carrying LoxP sites flanking the three exons of the FGF21 gene with albumin-Cre mice on the same genetic background. We confirmed the specific deletion of *Fgf21* in livers of adult *Fgf21^hep-/-^* mice indicating that most hepatic FGF21 expression is from hepatocytes **(Fig. 1A,B)**. As prior studies have shown that FGF21 reduces sweet preference (Talukdar *et al*., 2016; Von Holstein-Rathlou *et al*., 2016; Iroz *et al*., 2017), we investigated whether the absence of hepatocyte FGF21 affects sucrose intake using the two-bottle preference assay (10% sucrose in drinking water *versus* water). Water consumption, which was measured daily for 3 days, confirmed that *Fgf21^hep-/-^*mice consumed more sweetened water than *Fgf21^hep+/+^* mice **(Fig. 1C).**

**Figure 1.**
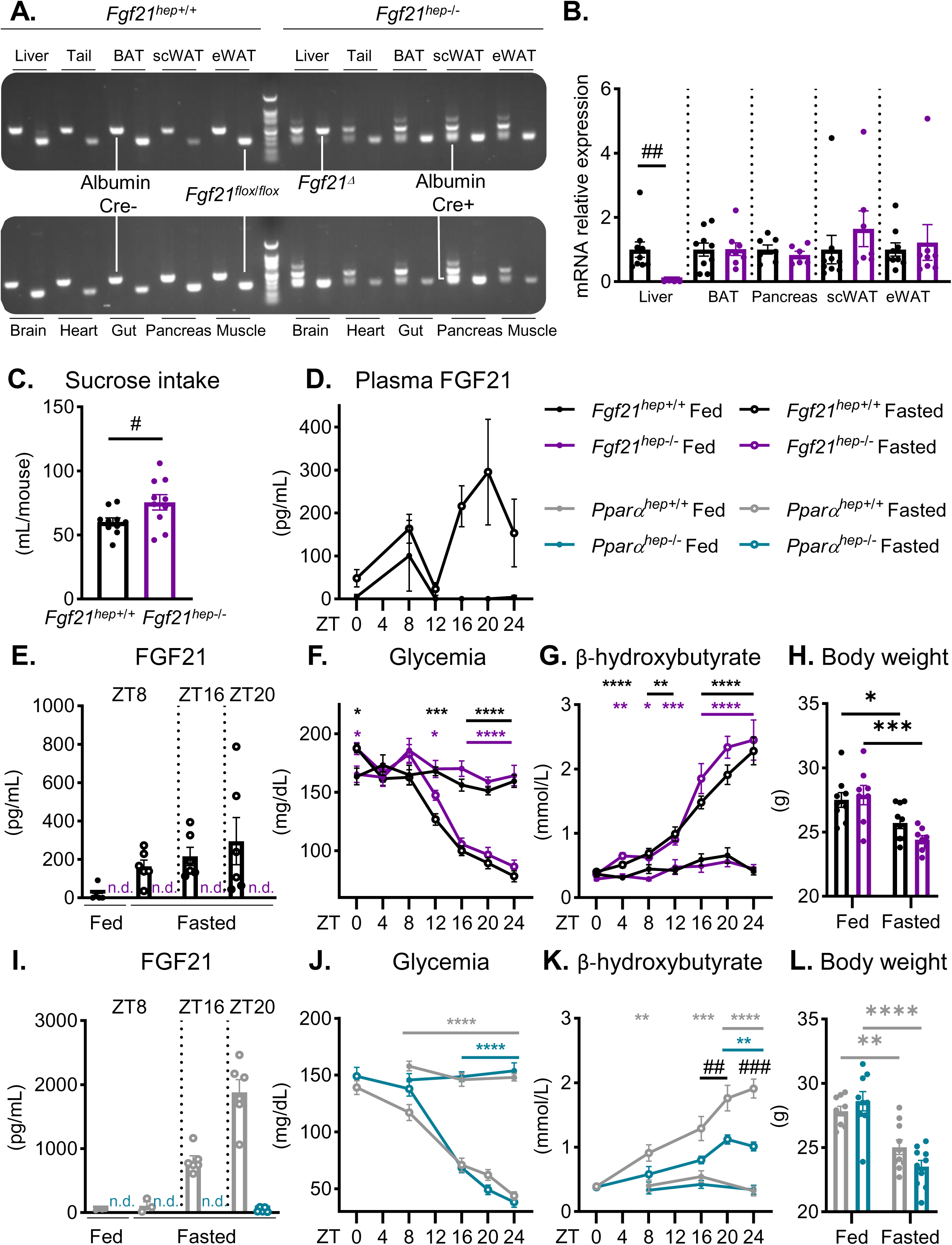
Characterisation of the hepatocyte-specific fibroblast growth factor 21 (FGF21) knockout mouse model. **(A)** PCR analysis of *Fgf21* floxed (*Fgf21^hep+/+^*) and *Albumin-Cre* (*Albumin-Cre^+/-^*) genes from male mice that are liver floxed (*Fgf21^hep+/+^*) or liver knockout (*Fgf21^hep-/-^)* for *Fgf21* using DNA extracted from different organs. **(B)** mRNA relative expression of *Fgf21* in liver, brown adipose tissue (BAT), pancreas, subcutaneous white adipose tissue (scWAT), epidemial white adipose tissue (eWAT) samples of *Fgf21^hep+/+^* and *Fgf21^hep-/-^* male mice measured by qRT-PCR (n=6-9 mice/group). **(C)** Sucrose intake during two bottle choice of 10% sucrose *vs* water for 4 days, in *Fgf21* liver floxed (*Fgf21^hep+/+^*) or liver knockout (*Fgf21^hep-/-^)* male mice (n=10 mice/group). **(D)** Liver floxed (*Fgf21^hep+/+^*) for *Fgf21* male mice were fed *ad libitum* or fasted for 24h and blood was collected at ZT0, ZT8, ZT12, ZT16, ZT20 and ZT24. FGF21 plasma level was determined by ELISA (n=6 mice/group). **(E-H)** *Fgf21* liver floxed (*Fgf21^hep+/+^*) or liver knockout (*Fgf21^hep-/-^)* male mice were fed *ad libitum* or fasted for 24h and blood was collected at ZT8, ZT16, ZT20. FGF21 plasma level was determined by ELISA (n=5-6 mice/group) **(E)**, blood glucose **(F)** and β-hydroxybutyrate levels **(G)** were monitored from ZT0 to ZT24, and body weight was measured after 24h fasting **(H)** (n=8-10 mice/group). **(I-L)** *Pparα* liver floxed (*Pparα^hep+/+^*) or *Pparα* liver knockout (*Pparα^hep-/-^*) mice were fed *ad libitum* or fasted for 24h and blood was collected at ZT8, ZT16, ZT20. FGF21 plasma level was determined by ELISA (n=5-6 mice/group) **(I)**, blood glucose **(J)** and β-hydroxybutyrate levels **(K)** were monitored from ZT0 to ZT24, and body weight was measured after 24h fasting **(L)** (n=9-10 mice/group). Data are presented as means±SEM. Significance of qPCR and sucrose intake datas are based on unpaired t-test. Significance of glycemia, ketomenia and body weight datas are based on 2-way ANOVA followed by Šídák’s multiple comparisons test * or # p<0.05; ** or ## p<0.01; *** or ### p<0.001; **** p<0.0001. * Fasting effect. # Genotype effect (C) or Genotype effect between fasting groups (K). (n.d.: not detected; ZT: Zeitgeber Time).

We next assessed the kinetics of plasma FGF21 levels in *Fgf21^hep+/+^* mice fed *ad libitum* or fasted for 24 hours. In agreement with previous reports (Montagner *et al*., 2016), we observed a peak of plasma protein expression at ZT8 in both fed and fasted groups during the light phase, corresponding to the sleep period of the animals. During the dark phase, plasma FGF21 concentrations increased and peaked at ZT20 only in fasted mice **(Fig. 1D)**. Circulating FGF21 was not detected in *Fgf21^hep-/-^* mice, confirming that the liver is the main source of plasma FGF21 (Markan *et al*., 2014) **(Fig. 1E)**. The decrease in fasting blood glucose levels and the concomitant increase in ketone bodies starting from ZT12 were similar between *Fgf21^hep+/+^* and *Fgf21^hep-/-^* mice **(Fig. 1F,G)**. The body weight loss was also comparable between genotypes after 24 hours of fasting **(Fig. 1H)**. Similar results were observed in fasted *Fgf21^hep+/+^* and *Fgf21^hep-/-^*females **(Fig. EV1A-D)**.

We also studied the effect of FGF21 deletion during a ketogenic diet, a condition that markedly induces FGF21 in the liver (Badman *et al*., 2007) **(Fig. EV1E)**. No significant difference in blood glucose, plasma ketone body levels, and body weight was observed between *Fgf21^hep+/+^*and *Fgf21^hep-/-^* mice fed a ketogenic diet for 9 days **(Fig. EV1F-H)**.

Liver FGF21 induction in response to starvation is highly controlled by the nuclear receptor PPARα in response to adipose tissue lipolysis (Badman *et al*., 2007; Inagaki *et al*., 2007; Montagner *et al*., 2016; Fougerat *et al*., 2022). Using mice with hepatocyte-specific deletion of *Pparα* (Montagner *et al*., 2016), we confirmed that fasting-induced FGF21 production was suppressed in the absence of PPARα in hepatocytes in both fed and fasted states **(Fig. 1I)**. As expected, glycemia over a 24h fasting challenge was unaffected by the absence of PPARα whereas fasted hepatocyte *Pparα*-decient mice (*Pparα^hep-/-^*) showed lower levels of plasma ketone bodies compared to fasted *Pparα^hep+/+^* mice **(Fig. 1J,K)** (Régnier *et al*., 2018). This observation suggests that fasting ketogenesis depends on PPARα, but not on FGF21. Body weight was decreased in response to fasting in both genotypes **(Fig. 1L)**.

These data confirmed that, during fasting, circulating FGF21 primarily originates from hepatocytes and is dependent on PPARα. However, *Fgf21* deletion in hepatocytes does not alter key metabolic adaptations to fasting such as ketogenesis, hypoglycemia, or weight loss. This observation contrasts with previous studies, which have shown that FGF21 is essential for ketogenesis (Badman *et al*., 2007; Inagaki *et al*., 2007; Potthoff *et al*., 2009). However, others reported no alteration in plasma ketone body levels in either whole-body FGF21 knockout mice or FGF21-overexpressing models during fasting (Inagaki *et al*., 2007; Badman *et al*., 2009; Hotta *et al*., 2009). Our findings in mice with hepatocyte-specific deletion of *Fgf21* establish that liver-derived FGF21 is dispensable for key metabolic responses to fasting including hypoglycemia and hepatic ketone body production.

### 2. Hepatocyte FGF21 is dispensable for fasting-induced responses in the liver

Fasting induces a major shift in hepatic metabolic homeostasis (Fougerat *et al*., 2024; Ruppert and Kersten, 2024). To investigate the role of endogenous hepatocyte FGF21 in fasting-induced changes in hepatic homeostasis, *Fgf21^hep+/+^* and *Fgf21^hep-/-^* mice were either fed *ad libitum* or fasted for 20 hours. Liver weight was significantly lower in fasted mice compared to fed animals but did not differ between the genotypes **(Fig. 2A)**. Hepatic triglyceride content and plasma alanine aminotransferase levels were also not affected by *Fgf21* deletion both in fed and fasted mice **(Fig. 2B,C)**. Consistent with these findings, hematoxylin and eosin staining of liver tissue did not reveal any difference between *Fgf21^hep+/+^* and *Fgf21^hep-/-^* mice **(Fig. 2D)**. Similarly, the deletion of *Fgf21* in hepatocytes did not affect liver weight and circulating alanine aminotransferase in fasted females **(Fig. EV2A,B)** nor in male mice fed a ketogenic diet **(Fig. EV2C,D)**.

**Figure 2.**
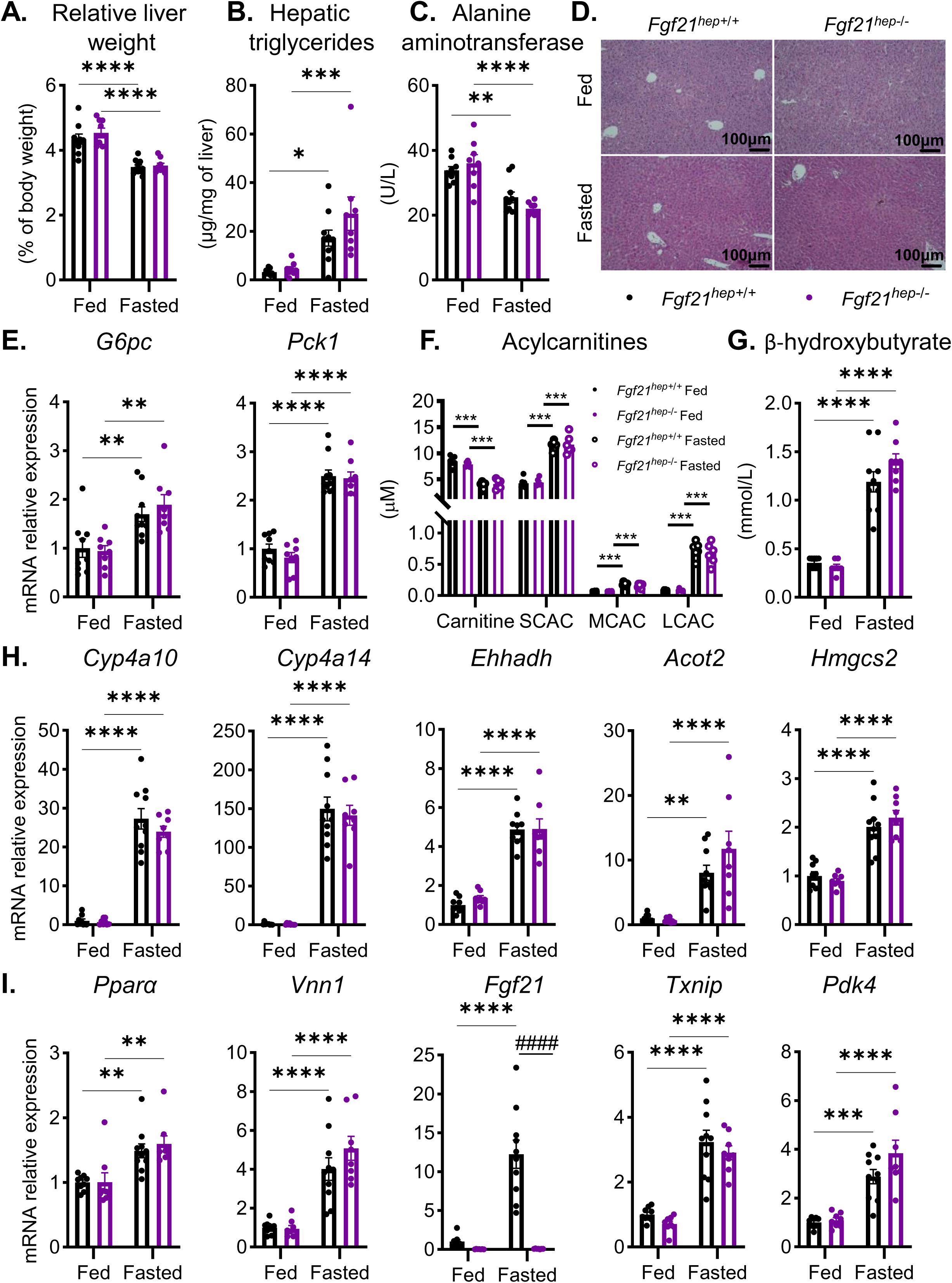
Hepatocyte-specific deletion of *Fgf21* does not affect fasting-induced hepatic responses. *Fgf21* liver floxed (*Fgf21^hep+/+^*) or *Fgf21* liver knockout (*Fgf21^hep-/-^*) mice were fed *ad libitum* or fasted for 20 hours. **(A)** Relative liver weight. **(B)** Hepatic triglyceride levels. **(C)** Plasma alanine aminotransferase levels. **(D)** Representative pictures of H&E staining of liver sections. Scale bars, 100µm. **(E)** mRNA relative expression of *G6pc and Pck1* in liver samples measured by qRT-PCR. **(F)** Plasma levels of carnitine, Short Chain Acylcarnitines (SCAC, C2-C5), Medium Chain Acylcarnitines (MCAC, C6-C12), and Long Chain Acylcarnitines (LCAC, C14-C18). **(G)** Plasma level of β-hydroxybutyrate. **(H)** mRNA relative expression of *Cyp4a10, Cyp4a14, Ehhadh, Acot2, Hmgcs2* and **(I)** *Pparα, Vnn1, Fgf21, Txnip, Pdk4* in liver samples measured by qRT-PCR. Data are presented as means±SEM from 8-10 mice/group (A-E,G-I) and from 6 mice/group (F). Significance are based on 2-way ANOVA followed by Šídák’s multiple comparisons test * p<0.05; ** p<0.01; *** p<0.001; **** or #### p<0.0001. * Fasting effect. # Genotype effect.

During fasting, the liver ensures the production of glucose and ketone bodies to supply peripheral tissues. After glycogen stores are depleted, glucose production depends on gluconeogenesis (Fougerat *et al*., 2024). In contrast with whole-body *Fgf21^-/-^* mice (Potthoff *et al*., 2009; Liang *et al*., 2014), we found that hepatic expression of key enzymes involved in gluconeogenesis, *G6pc* and *Pck1*, was increased in response to fasting in both *Fgf21^hep+/+^*and *Fgf21^hep-/-^* mice **(Fig. 2E).** These findings were further supported by a similar glucose production in the two genotypes, as assessed by the pyruvate tolerance test indicating that hepatocyte *Fgf21* deletion did not affect fasting-induced hepatic gluconeogenesis **(Fig. EV3)**.

If fasting persists, gluconeogenesis decreases to prevent muscle protein breakdown while fatty acids derived from adipose tissue lipolysis are utilized to produce ketone bodies in the liver. Once in the liver, fatty acids are converted into acylcarnitines which, in turn, are beta-oxidized in mitochondria (Fougerat *et al*., 2024). In line with this, we found that fasting decreased the plasma level of free carnitine and increased short-, medium-, and long-chain acylcarnitine levels with no significant difference between genotypes **(Fig. 2F)**. The increase in plasma ketone bodies and hepatic expression of genes involved in the regulation of fatty acid catabolism (β-oxidation and ketogenesis) was also similar in *Fgf21^hep+/+^* and *Fgf21^hep-/-^* mice in response to fasting **(Fig. 2G,H)**. The induction of fatty acid oxidation and ketogenesis during fasting is mainly orchestrated transcriptionally by the nuclear receptor PPARα. Given that hepatic PPARα activity is regulated by adipose tissue lipolysis (Fougerat *et al*., 2022) and that FGF21 has been reported to regulate lipolysis (Inagaki *et al*., 2007; Hotta *et al*., 2009; Chen *et al*., 2011), FGF21 may exert a negative feedback or stimulate PPARα activity in hepatocytes during fasting. Except that of *Fgf21*, the expression of well-known PPARα target genes during fasting was unchanged in the absence of FGF21 in hepatocytes (Régnier *et al*., 2018) **(Fig. 2I)**. Expression of these genes was also not impacted by *Fgf21* deletion in fasted females and in males fed a ketogenic diet **(Fig. EV2E-H)**. Similar profiles were observed for genes involved in hepatic autophagy, a conserved catabolic process activated in the liver to recycle intracellular substrates and remove dysfunctional proteins during fasting **(Fig. EV4)**.

Together, these data indicate that hepatocyte FGF21 does not contribute to fasting-induced key hepatic metabolic pathways such as gluconeogenesis, fatty acid β-oxidation and ketogenesis. Additionally, our results do not support a role for hepatocyte FGF21 in the PPARα-dependent responses in the liver.

### 3. Hepatocyte FGF21 deletion does not influence liver gene expression and metabolites in response to fasting

Fasting induces a major change in liver gene expression and metabolome (Goldstein and Hager, 2015; Ruppert and Kersten, 2024). To further investigate the contribution of FGF21 on liver gene expression, we performed gene expression profiling on livers of fed and fasted *Fgf21^hep+/+^* and *Fgf21^hep-/-^* mice. Principal-Component Analysis (PCA) showed that differences in gene expression were mostly observed between fed and fasted mice and indicated minimal effects of *Fgf21* deletion on the hepatic transcriptome **(Fig. 3A)**. Volcano plot comparing differentially expressed genes (DEGs) between fed and fasted conditions revealed that *Fgf21* was the only gene significantly upregulated in response to fasting in *Fgf21^hep+/+^* compared to *Fgf21^hep−/−^* mice **(Fig. EV5A)**. Hierarchical clustering on the DEGs (Fold-Change (FC) > 1.5; p_adj_ < 0.05) confirmed the marked discrimination between fed and fasted mice, and identified two clusters showing specific gene expression in response to fasting, which were not dependent on *Fgf21* expression in hepatocytes **(Fig. 3B)**. In the first cluster, 4519 genes were upregulated in response to fasting **(Fig. 3C)**. Gene Ontology (GO) analysis revealed that these genes were most significantly associated with catabolic metabolic processes, mostly PPARα targets, as identified through transcription factor enrichment **(Fig. 3D,E)**. Consistent with these observations, the most upregulated genes by fasting in *Fgf21^hep+/+^* mice were known PPARα target genes. These genes were also upregulated in *Fgf21^hep−/−^* mice, except for the deleted *Fgf21* **(Fig. EV5B)**. The 4549 genes in cluster 2 exhibited lower mRNA expression in fasted mice compared to fed mice **(Fig. 3B,C)**. This cluster was most significantly associated with ‘steroid biosynthetic process’ and enriched in SREBF1, SP1, and MLXIP target genes **(Fig. 3D,E)**. The most significantly affected genes in cluster 2 included genes involved in lipogenesis and cholesterol metabolism, such as *Fasn*, *Pnpla5*, *Pcsk9*, and *Sqle* **(Fig. EV5C)**. Next, we plotted the FC values for *Fgf21^hep−/−^* mice in response to fasting (x-axis) against those for *Fgf21^hep+/+^* mice under the same conditions (y-axis). A 45-degree trend line was observed, confirming that fasting exerts similar effects on liver gene expression in *Fgf21^hep+/+^* and *Fgf21^hep−/−^* mice **(Fig. 3F)**. Consistent with our previous findings **(Fig. 2H,I)**, fasting-induced PPARα-sensitive genes (Régnier *et al*., 2018) clustered closely along the trend line, with the exception of *Fgf21*. Finally, we performed a Pearson correlation analysis to identify genes whose expression levels are correlated with that of *Fgf21*, using a correlation coefficient threshold > 0.7. The expression profiles of these genes were not significantly different between fasted *Fgf21^hep+/+^*and fasted *Fgf21^hep-/-^* mice **(Fig. EV5D)**.

**Figure 3.**
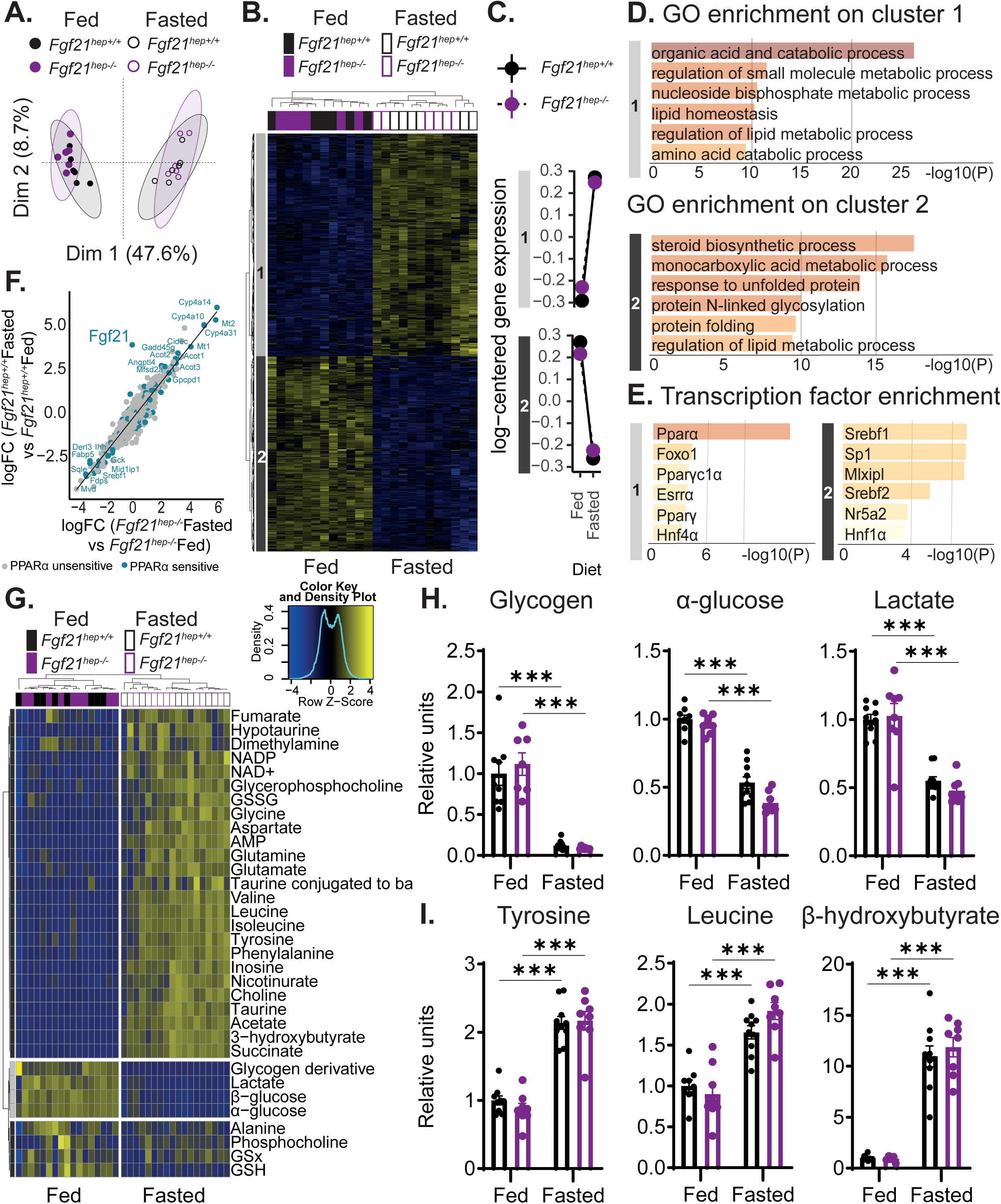
Hepatocyte FGF21 deletion does not influence liver gene expression and metabolites in response to fasting. *Fgf21* liver floxed (*Fgf21^hep+/+^*) or *Fgf21* liver knockout (*Fgf21^hep-/-^*) mice were fed *ad libitum* or fasted for 20 hours. **(A)** Principal component analysis (PCA) score plots of whole-liver transcriptome dataset. **(B)** Microarray experiment performed with liver samples. Hierarchical clustering shows the definition of two gene clusters (FC > 1.5; p_adj_< 0.05). **(C)** Representation of the mean cluster profiles in each cluster. **(D-E)** Gene Ontology (GO) analysis in heatmap cluster 1 and cluster 2 **(D)** and enrichment of transcription factors **(E)**. **(F)** Representation of fasting hepatic PPARα-dependent genes on the correlation of fasting-dependent genes in *Fgf21* liver floxed (*Fgf21^hep+/+^*) mice vs. fasting-dependent genes in *Fgf21* liver knockout (*Fgf21^hep-/-^*) mice. **(G)** Heatmap of liver metabolomic dataset. Hierarchical clustering shows the definition of three metabolite clusters. **(H-I)** Area under the curve of the ^1^H-NMR spectra for representative metabolites that are significantly repressed **(H)** and induced **(I)** by fasting. Data are presented as means±SEM from 6 mice/group (A-F) and from 8-10 mice/group (G-I). Significance are based on 2-way ANOVA followed by Šídák’s multiple comparisons test. *** p<0.001. * Fasting effect.

To further explore the potential impact of *Fgf21* deficiency, we next performed hepatic metabolomic profiling of aqueous metabolites using ^1^H-NMR. This analysis confirmed the presence of marked fasting-dependent hepatic metabolites which remained independent of *Fgf21* expression in hepatocytes **(Fig. 3G)**. Among the metabolites reduced during fasting, we found glycogen, α-glucose, and lactate, which are utilized by the liver as energy substrates **(Fig. 3H)**. Conversely, the concentrations of several amino acids were increased in fasted mice, likely deriving from protein degradation. Hepatic β-hydroxybutyrate, the most abundant ketone body, was also increased by fasting in both genotypes **(Fig. 3I)**.

Collectively, these data indicate that the fasting-induced changes in hepatic gene expression and metabolome are not dependent on hepatocyte FGF21. Given the significant increase in circulating FGF21 levels during fasting in *Fgf21^hep+/+^* mice, these results were somewhat unexpected. We confirmed that hepatic expression of the FGF21 receptor complex, including *Fgfr1c* and *β-Klotho*, was similar in *Fgf21^hep+/+^* and *Fgf21^hep−/−^* mice in both fed and fasted mice **(Fig. EV5E).** In addition, hepatic expression of several other hepatokines remained unaltered, ruling out compensatory upregulation mechanisms in the absence of FGF21 **(Fig. EV6)**.

### 4. Hepatocyte FGF21 deletion does not influence adipose tissue responses to fasting

Having established that autocrine FGF21 signaling is not essential for fasting-induced responses in the liver, we next asked whether hepatic FGF21 influences adipose tissue function. While it is clear that FGF21 signaling in adipose tissue has insulin-sensitizing (Arner *et al*., 2008; Li *et al*., 2009) and lipolysis inhibitory effects in obese mice (Ding *et al*., 2012; BonDurant *et al*., 2017), the contribution of FGF21 to adipose tissue lipolysis during fasting remains debated. Previous studies have reported that global deletion of *Fgf21* increases fasting-induced lipolysis (Hotta *et al*., 2009; Chen *et al*., 2011) whereas others have shown that circulating free fatty acid levels are not affected by the absence of FGF21 (Badman *et al*., 2009; Potthoff *et al*., 2009). More recent work demonstrated that acute deletion of hepatic FGF21 does not affect adipose tissue lipolysis in fasted mice (Sostre-Colón *et al*., 2022). We therefore investigated the potential contribution of hepatocyte-derived FGF21 to the physiology of the adipose tissues.

We first measured the expression of *Fgf21* and its receptor complex in subcutaneous (sc) and epididymal (ep) white adipose tissues (WAT) and in brown adipose tissue (BAT), which was not affected by hepatocyte *Fgf21* deletion in the 3 tissues **(Fig. EV7A-C)**. The relative weight of WATs and BAT was not significantly different between groups **(Fig. 4A,B and EV8A)**. Circulating free fatty acids and glycerol were increased in fasted mice compared to fed mice but not different between the two genotypes, indicating that the lack of FGF21 in hepatocytes does not influence WAT lipolysis during fasting **(Fig. 4C,D)**. Similar results were observed in males fed a ketogenic diet as well as in fasted females **(Fig. EV9A-H)**.

**Figure 4.**
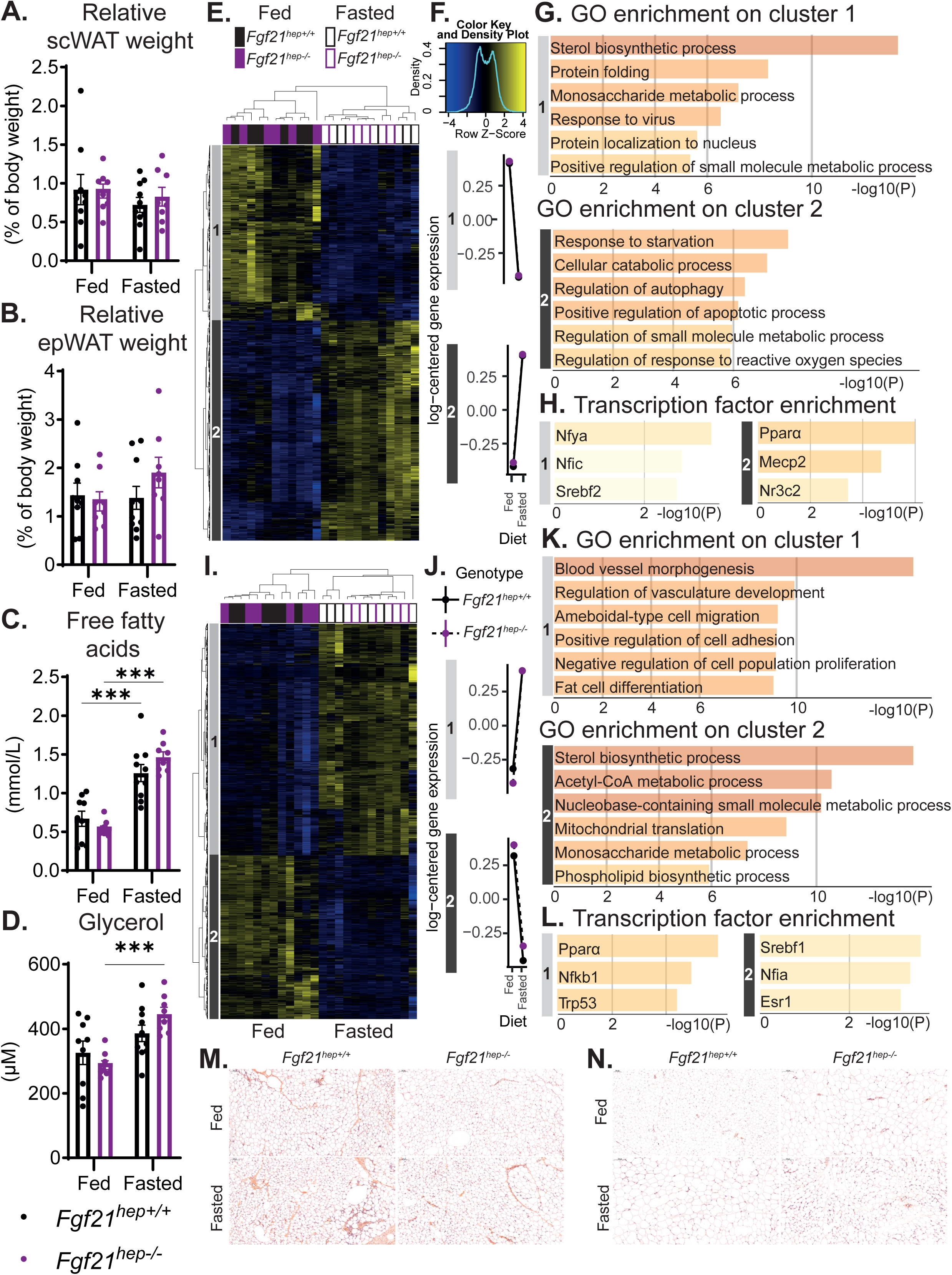
Hepatocyte FGF21 deletion does not influence adipose tissue responses to fasting. *Fgf21* liver floxed (*Fgf21^hep+/+^*) or *Fgf21* liver knockout (*Fgf21^hep-/-^*) mice were fed *ad libitum* or fasted for 20 hours. **(A)** Relative sub-cutaneous white adipose tissue (scWAT) weight. **(B)** Relative epidemial white adipose tissue (epWAT) weight. **(C)** Plasma free fatty acids. **(D)** Plasma level of glycerol. **(E, I)** Microarrays experiments performed with scWAT **(E)** and epWAT **(I)** samples. For each heatmap, hierarchical clustering shows the definition of two gene clusters (FC > 1.5; p_adj_< 0.05). **(F, J)** Representation of the mean cluster profiles in each cluster for scWAT **(F)** and epWAT **(J)** dataset. **(G, K, H, L)** Gene Ontology (GO) analysis in heatmap cluster 1 and cluster 2 of scWAT **(G)** and epWAT dataset **(K)** and their respective enrichment of transcription factors **(H, L)**. **(M-N)** Representative pictures of H&E staining of scWAT **(M)** and epWAT **(N)** sections. Scale bars, 100μm. Data are presented as means±SEM from 8-10 mice/group (A-D) and from 6 mice/group (E-L). Significance are based on 2-way ANOVA followed by Šídák’s multiple comparisons test. *** p<0.0001. * Fasting effect.

To further examine the potential effect of hepatocyte *Fgf21* deletion on adipose tissues, we performed microarray analysis on scWAT, epWAT and BAT gene expression. In the three tissues, hierarchical clustering using the DEGs (FC > 1.5; p_adj_< 0.05) identified two main gene clusters and revealed a marked discrimination only between fed and fasted animals, independent of *Fgf21* expression in hepatocytes **(Fig. 4E,I and EV8B)**. Volcano plots comparing DEGs between fasted and fed conditions in both genotypes further supported this conclusion. Notably, no gene met the significant p_value threshold **(Fig. EV7D-F)**. The most up- and down-regulated genes in response to fasting in scWAT, epWAT, and BAT are shown in **Fig. EV7G-I**. In scWAT, cluster 1 comprised genes that are down-regulated following fasting. These genes are mainly associated with “sterol biosynthetic process”. In contrast, genes in cluster 2 exhibited a higher expression in fasted mice and were involved in “response to starvation”, “cellular catabolic process”, and “regulation of autophagy” **(Fig. 4F,G)**. In epWAT, genes in cluster 1 were up-regulated by fasting and associated with “blood vessel morphogenesis”, “regulation of vasculature development”, and “fat cell differentiation”, indicating transcriptional changes related to tissue remodeling and cellular plasticity. As for scWAT, genes with lower expression upon fasting (cluster 2) were mainly involved in “sterol biosynthetic process” **(Fig. 4J,K)**. Similarly, genes with lower expression in fasted BAT were involved in the “sterol biosynthetic process” while genes up-regulated by fasting were associated with “cellular response to starvation” **(Fig. EV8C,D)**. In all three adipose tissues, genes induced by fasting were identified mostly as PPARα target genes while SREBP1 and SREBP2 targets are repressed by fasting **(Fig. 4H, L and EV8E)**. Histological analysis of WATs and BAT sections stained with hematoxylin and eosin (H&E) revealed no marked difference in adipocyte morphology **(Fig. 4M,N and EV8F)**.

Taken together, these results provide a detailed characterization of gene expression in three different adipose tissues in response to fasting. In line with prior studies, we found that fasting induces major changes in adipose tissue gene expression (Ruppert and Kersten, 2025). The regulation of adipose tissue metabolism during fasting is mainly driven by changes in the plasma levels of several hormones, such as reduction in plasma insulin, that mediate transcriptional and posttranslational regulations of key enzymes involved in pathways aimed at storing energy (extracellular lipolysis, triglyceride synthesis) and liberating fatty acids (intracellular lipolysis) (Kersten, 2023; Ruppert and Kersten, 2025). Our findings indicate that these adipose tissue responses to fasting are largely independent of *Fgf21* expression in hepatocytes and circulating FGF21 level.

### 5. Hepatocyte FGF21 drives protein preference following fasting

Beyond its role in metabolic regulation, FGF21 has been implicated in the control of nutrient-driven behaviors, particularly in protein appetite and sweet preference (Laeger *et al*., 2014; Von Holstein-Rathlou *et al*., 2016; Hill *et al*., 2019; Solon-Biet *et al*., 2023; Kim *et al*., 2024; Nicolaisen *et al*., 2025). To explore this aspect, we examined whether the deletion of *Fgf21* in hepatocytes may affect systemic energy expenditure and macronutrient preference.

We first performed metabolic phenotyping *via* indirect calorimetry in fed, fasted and refed *Fgf21^hep+/+^* and *Fgf21^hep−/−^* mice. There was no difference in physical activity between genotypes under all nutritional conditions **(Fig. EV10A)**. In the fed state, oxygen consumption (VO_2_), carbon dioxide production (VCO₂), and total energy expenditure (EE) were slightly higher in *Fgf21^hep−/−^* mice than in *Fgf21^hep+/+^* mice at the beginning of the active dark period **(Fig. EV10B-D)**. However, the expiratory exchange ratio was similar in both genotypes **(Fig. EV10E)**. None of these parameters was strongly affected by *Fgf21* deletion in the fasted state **(Fig. EV10B-E)**. Most changes were observed during refeeding with *Fgf21^hep−/−^* mice exhibiting higher VO_2_, VCO₂, and EE than *Fgf21^hep+/+^* mice **(Fig. EV10B-D)**. As for fed mice, the expiratory exchange ratio was similar in the two genotypes during refeeding, indicating similar substrate utilization **(Fig. EV10E)**.

We then investigated the impact of hepatocyte *Fgf21* deletion during refeeding following a fasting period. Food choice after fasting represents a critical nutritional checkpoint during which animals must prioritize specific macronutrients to restore metabolic balance. Protein is unique among the macronutrients as it provides essential amino acids. A reduction in protein intake triggers adaptive physiological and behavioral changes in food intake and preference to prevent protein deficiency (Kim *et al*., 2024). FGF21 is robustly induced in the liver in response to protein restriction (De Sousa-Coelho, Marrero and Haro, 2012; Laeger *et al*., 2014; Solon-Biet *et al*., 2023; Nicolaisen *et al*., 2025) and acts in the brain to enhance protein appetite while suppressing sugar preference (Laeger *et al*., 2014, 2016; Talukdar *et al*., 2016; Hill *et al*., 2019).

To test whether hepatic FGF21 contributes to nutrient selection following fasting, we performed a two-choice preference test after fasting in *Fgf21^hep+/+^* and in *Fgf21^hep−/−^* mice. During the baseline acclimation phase, no significant difference in 24-hour food intake was observed between genotypes **(Fig. EV11A).** Food intake during 24-hour refeeding after 20 hours of fasting was also comparable between genotypes **(Fig. EV11B)**. In a first two-food choice test, mice were fasted for 20 hours and then exposed simultaneously *ad libitum* to a low-protein diet (6.5% protein, 80.4% carbohydrate, 3.8 kcal/g) and to a high-protein diet (42.6% protein, 44.3% carbohydrate, 3.8 kcal/g) for 24 hours **(Table 1 and Fig. 5A).** *Fgf21^hep+/+^* mice displayed a balanced consumption of both diets. In contrast, *Fgf21^hep−/−^* mice consumed significantly less of the high-protein diet than low-protein diet **(Fig. 5B)**. Preference analysis confirmed that *Fgf21^hep+/+^*mice displayed a relative balance between diets (52.72 % ± 8.97 preference high/low), while *Fgf21^hep−/−^* mice showed an impaired ability to shift toward the high-protein diet (8.18 % ± 4.58 preference high/low) **(Fig. 5C)**, from the first hours of refeeding **(Fig. EV11C)**.

**Figure 5.**
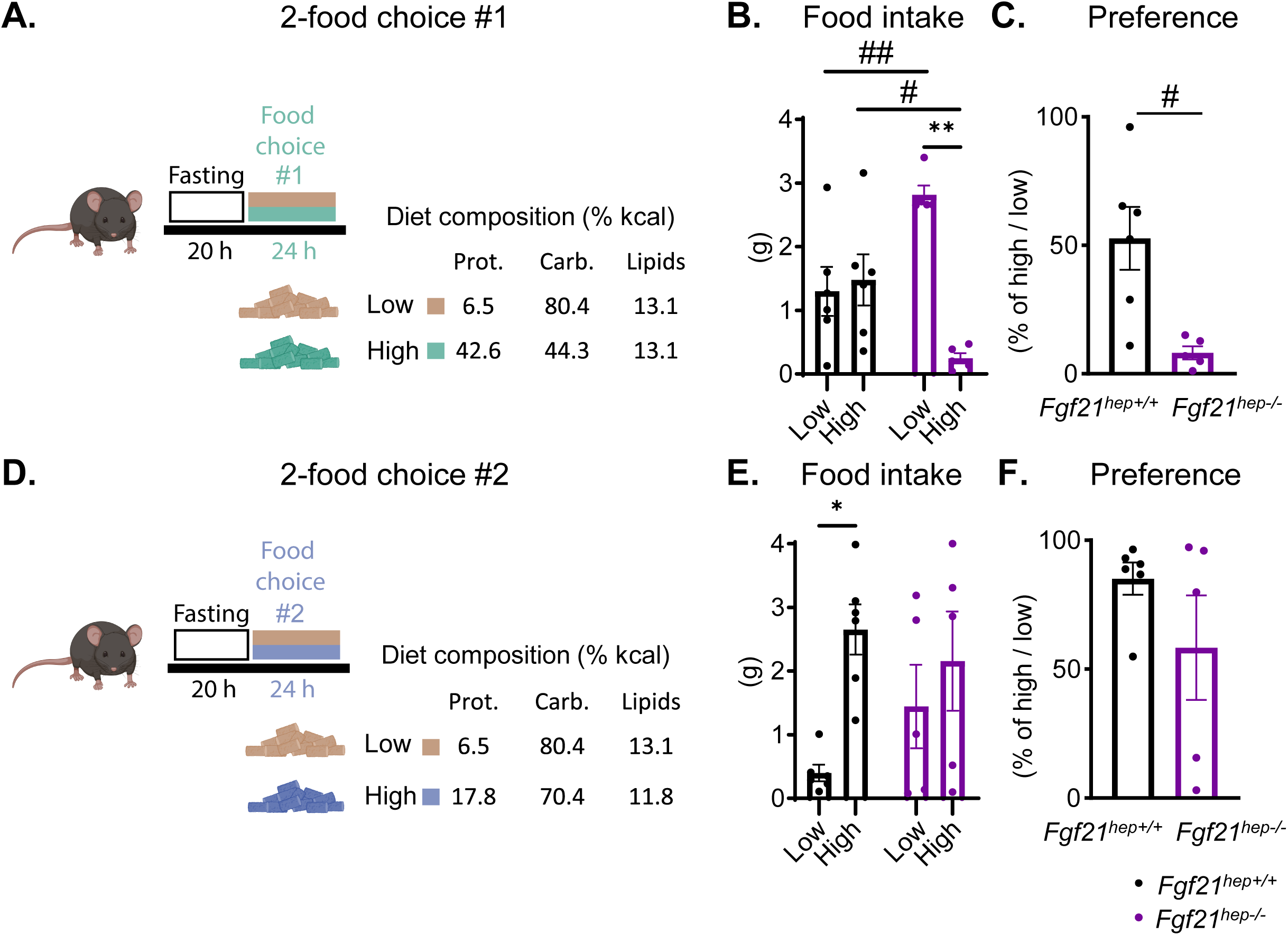
Hepatic FGF21 is required for protein preference after fasting. (A-C) *Fgf21* liver floxed (*Fgf21^hep+/+^*) or *Fgf21* liver knockout (*Fgf21^hep-/-^*) mice were fasted for 20 hours and were exposed *ad libitum* to a low-protein diet (low-protein/high-carb diet: 6.5% protein, 80.4% carbohydrate) and a high-protein diet (high-protein/low-carb diet: 42.6% protein, 44.3% carbohydrate) for 24 hours **(A)**. Food intake **(B)** and preference for the high-protein diet **(C)** over 24h (*P*-value of F-test for variance comparison: 0.0068). **(D-F)** *Fgf21* liver floxed (*Fgf21^hep+/+^*) or *Fgf21* liver knockout (*Fgf21^hep-/-^*) mice were fasted for 20 hours and were exposed *ad libitum* to a low-protein diet (low-protein/high-carb diet: 6.5% protein, 80.4% carbohydrate) and a high-protein diet (high-protein/high-carb diet: 17.8% protein, 70.4% carbohydrate) for 24 hours **(D)**. Food intake **(E)** and preference for the high-protein diet **(F)** over 24h (*P*-value of F-test for variance comparison: 0.0333). Mice were singly housed. The percentage of preference was calculated for each mouse by dividing the intake of low-protein diet or high-protein diet by the total food intake (low-protein + high-protein diets). Data are presented as means±SEM from 5-6 mice/group. Significance are based on 2-way repeated measures ANOVA or unpaired *t*-test with Welch’s correction for percentage of preference. * or # p<0.05; ** or ## p<0.001. * Food choice effect. # Genotype effect.

**Table 1:**
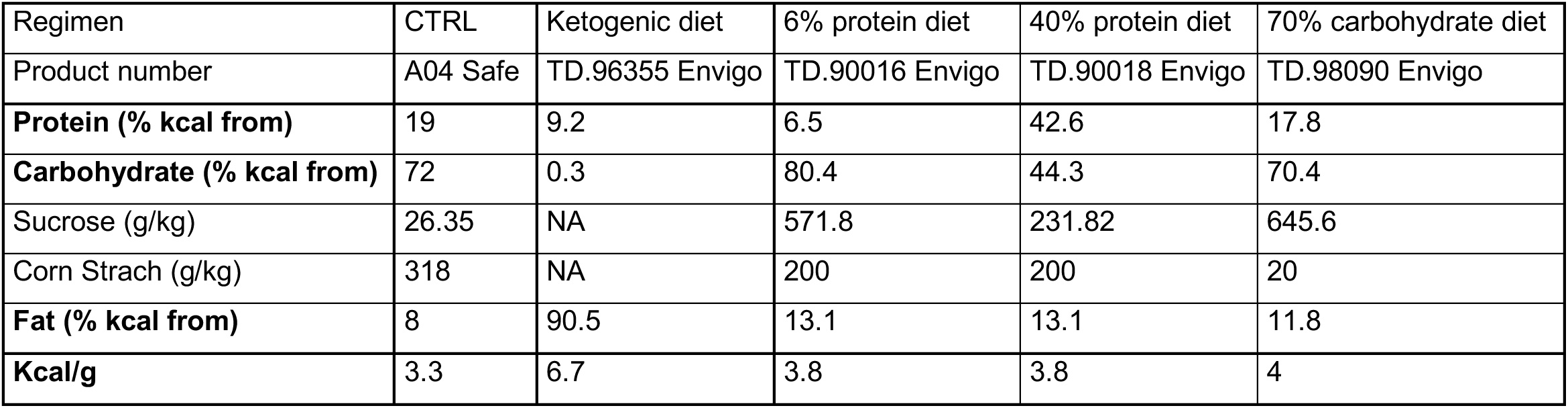
Diet composition.

In the first assay, the high-protein diet also had a lower carbohydrate content compared to the low-protein diet (44.3% *vs* 80.4%). To distinguish between protein and carbohydrate effects, we performed a second two-choice test in which both high- and low protein diets (17.8% and 6.5% protein, respectively) were adjusted with a comparable amount of carbohydrates (70.4% and 80.4% carbohydrate, respectively) **(Table 1 and Fig. 5D)**. In this second assay, *Fgf21^hep+/+^* mice displayed a marked preference for the high-protein diet straightaway (85.10 % ± 6.19 preference high/low), whereas *Fgf21^hep−/−^* mice consumed the same amount of both diets (58.35 % ± 17.53 preference high/low) **(Fig. 5E,F and EV11D)**. Importantly, total food intake was not different between genotypes **(Fig. 5B, E)**, indicating that FGF21 influences nutrient selection toward proteins rather than global caloric intake.

Prior studies also supported a key role of FGF21 in driving protein appetite under conditions of dietary protein restriction (Hill *et al*., 2019; Solon-Biet *et al*., 2023) and FGF21 signaling in the brain is required for this adaptive response (Hill *et al*., 2019). Pharmacological administration of FGF21 has also been reported to suppress sweet and alcohol preference (Dushay *et al*., 2015; Talukdar *et al*., 2016; Von Holstein-Rathlou *et al*., 2016; Flippo *et al*., 2022). Recent integrative models propose that this hormone coordinates multiple behavioral axes to adapt macronutrient intake to nutritional needs (Solon-Biet *et al*., 2023). Our findings strongly support this model and suggest a liver-brain endocrine axis to mediate preference for proteins following a fasting period. FGF21 signals through a receptor complex comprising a FGF receptor, primarily FGFR1c, and the co-receptor β-Klotho (KLB) (Lee *et al*., 2018), which are both expressed in specific regions of the brain, including the hypothalamus and the hindbrain (Bookout *et al*., 2013; Liang *et al*., 2014; Jensen-Cody *et al*., 2020). Additional studies will be required to precisely determine on which neurons liver-derived FGF21 acts to mediate its effect on protein preference following fasting. Recent data implicating hepatic FGF21 in sex-specific adaptations to juvenile protein malnutrition reinforce its role as a nutrient-sensitive endocrine coordinator (preprint : Joly et al., 2025). Conversely, a brain to liver axis regulated by FGF21 has been recently reported as a therapeutical target for metabolic liver disease. Pharmacological FGF21 administration in mice with diet-induced MASH decreased liver triglyceride content by acting on glutamatergic neurons in the central nervous system to increase sympathetic nerve activity to the liver (Rose *et al*., 2025).

In conclusion, our results confirm and extend previous findings (Markan *et al*., 2014; Montagner *et al*., 2016) showing that during fasting PPARα-dependent circulating FGF21 originates from hepatocytes. Unlike other studies in *Fgf21^-/-^* mice (Badman *et al*., 2009; Hotta *et al*., 2009; Potthoff *et al*., 2009; Liang *et al*., 2014), our work reveals that hepatocyte-restricted deletion of *Fgf21* does not influence fasting response. It does not exert any autocrine effect on key hepatic pathways such as gluconeogenesis or ketogenesis (Goldstein and Hager, 2015; Ruppert and Kersten, 2024). These data are in line with one report showing that acute AAV-mediated deletion of hepatocyte *Fgf21* does not influence basal glucose homeostasis or lipolysis in mice (Sostre-Colón *et al*., 2022).

Moreover, our combined genome and metabolome analysis suggest that FGF21 is dispensable for fasting-induced changes in liver homeostasis. The fasting response relies on a coordinated metabolic dialogue between adipose tissue and the liver (Jaeger *et al*., 2015; Fougerat *et al*., 2022). Previous studies have shown that recombinant FGF21 administration influences adipose tissues in mice (Inagaki *et al*., 2007; Coskun *et al*., 2008; Xu *et al*., 2009; Kliewer and Mangelsdorf, 2019). Moreover, hepatocyte-specific deletion of *Fgf21* alters liver to adipose tissue dialogue in mouse models of liver-specific insulin resistance (Sostre-Colón *et al*., 2021) and in diet-induced obesity (Vernia *et al*., 2014, 2016).

We therefore questioned whether hepatocyte-derived FGF21 production may influence adipose tissue homeostasis in the context of fasting. In line with previous reports (Ruppert and Kersten, 2025), we found that fasting induces major changes in WAT and BAT gene expression. However, these changes occur independently of hepatocyte FGF21. While we found that hepatocyte FGF21 is dispensable for metabolic adaptations during fasting, our results indicate that it becomes essential during the refeeding period to guide macronutrient selection toward proteins.

Altogether, our findings suggest a reconceptualization of liver FGF21 not as a direct regulator of fasting-induced control of metabolism, but rather as an endocrine signal that promotes protein appetite and the search for essential nutrients after food deprivation.

## Methods

### Reagents and Tools Table

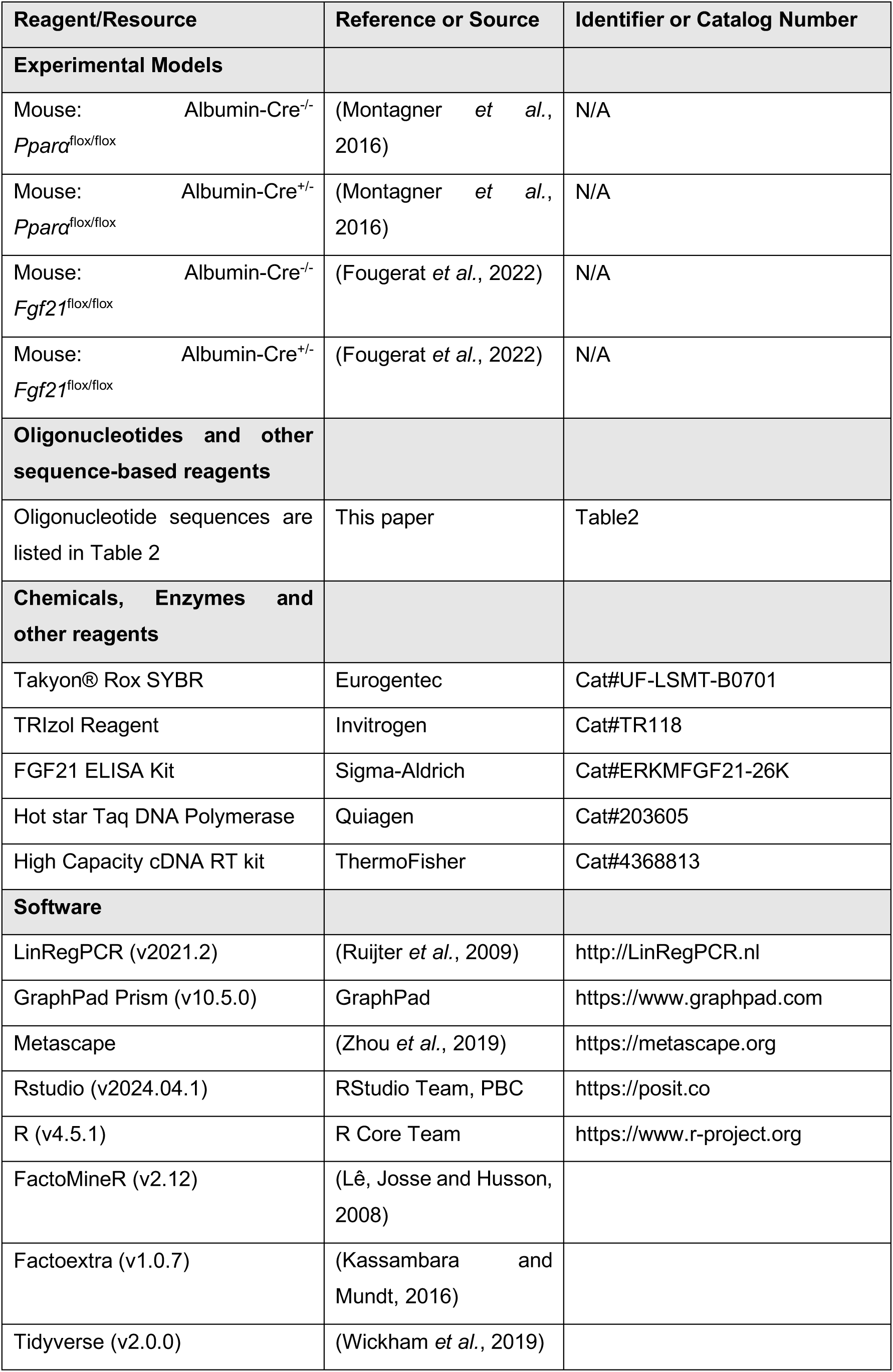
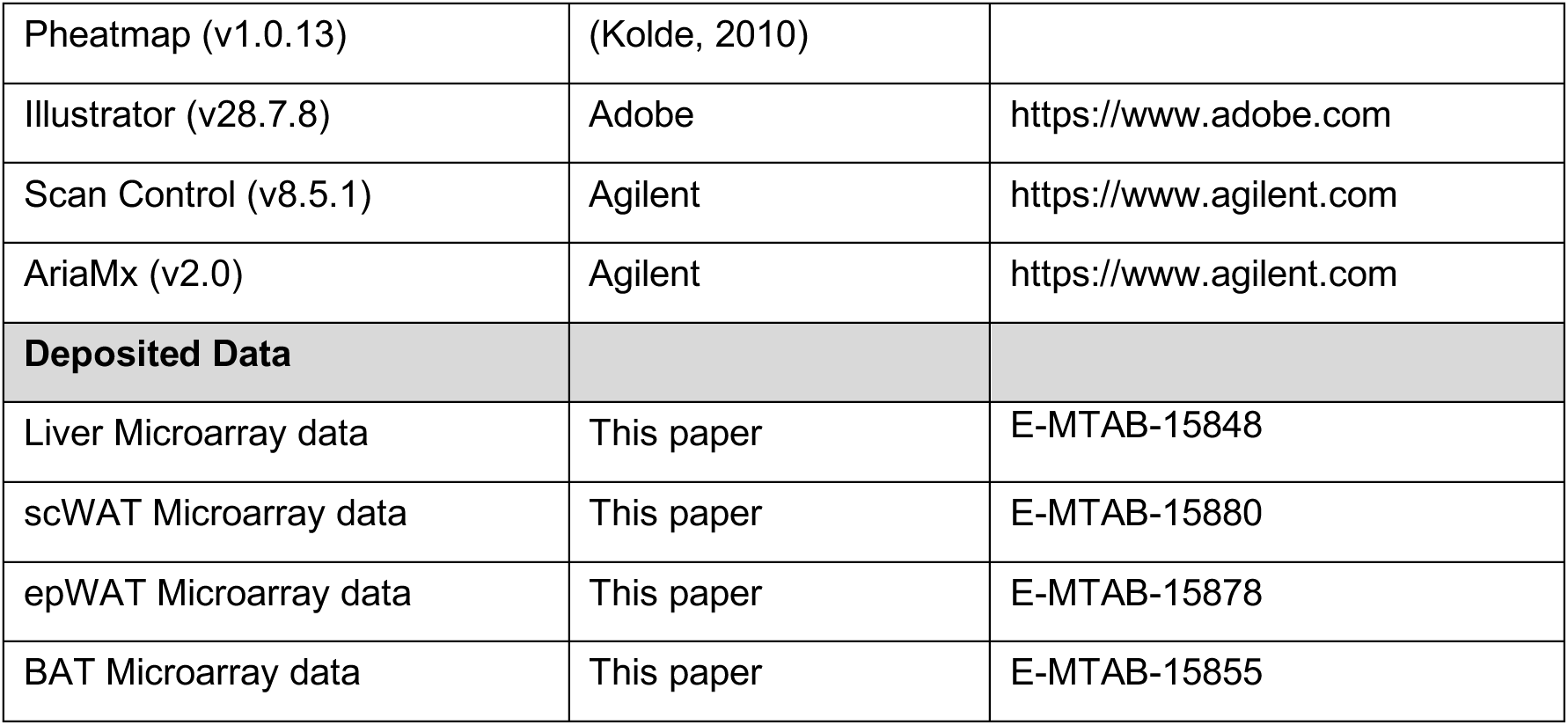

### Methods and Protocols

#### Mice

*In vivo* studies were performed in compliance with the European guidelines for the use and care of laboratory animals, and approved by an independent ethics committee under the authorization numbers 31741-2021052011598985. All mice were housed at 21–23°C on a 12-h light (ZT0–ZT12)/12-h dark (ZT12–ZT24) cycle and had free access to the standard rodent diet (Safe 04 U8220G10R) and tap water. The ketogenic diet (TD. 96355, INOTIV), the 6% protein diet (TD.90016, INOTIV), the 40% protein diet (TD.90018, INOTIV) and the 70% carbohydrate diet (TD.98090, INOTIV) were used for mouse dietary intervention. Their compositions were listed in **Table 1**.

ZT stands for Zeitgeber time; ZT0 is defined as the time when the lights are turned on. Mice used in this study were sacrificed at ZT20, unless stated otherwise. All experiments were performed in 10–13 week-old mice.

*Fgf21* hepatocyte-specific knockout (*Fgf21^hep-/-^*) mice were generated at INRAE’s rodent facility (Toulouse, France) by mating the floxed-*Fgf21* mouse strain (B6.129S6(SJL)Fgf21 < tm1.2Djm>/J provided by The Jackson Laboratory) with C57BL/6J albumin-Cre transgenic mice, to obtain albumin-Cre^+/-^*Fgf21*^flox/flox^ mice, as described previously (Fougerat *et al*., 2022). Albumin-Cre^-/-^*Fgf21*^flox/flox^ (*Fgf21^hep+/+^)* littermates were used as controls.

*Pparα* hepatocyte-specific knockout (*Pparα^hep-/-^*) mice were generated at INRAE’s rodent facility (Toulouse, France) by mating the floxed-*Pparα* mouse strain with C57BL/6J albumin-Cre transgenic mice, as described previously (Montagner *et al*., 2016), to obtain albumin-Cre^+/-^*Pparα*^flox/flox^ mice. Albumin-Cre^-/-^*Pparα*^flox/flox^ (*Pparα^hep+/+^*) littermates were used as controls.

#### Genomic DNA

Genomic DNA was extracted from liver, subcutaneous white adipose tissue (scWAT), epididymal white adipose tissue (epWAT), brown adipose tissue (BAT), heart, brain, pancreas, gut, and muscle samples, which were stored at -80°C following euthanasia. Tissue samples were homogenized in a solution containing 0.5M EDTA, 5M NaCl, 1M Tris-HCl (pH 7.4), 10% SDS, and 20mg/mL proteinase K (pH 8). The mixtures were incubated at 56°C with continuous shaking at 300 rpm for 3 hours. Following incubation, the samples were cooled on ice for 10 minutes and centrifuged. The supernatant was collected and mixed. For DNA extraction, 400µL of a chloroform:phenol:isoamyl alcohol (24:25:1) was added to the samples, which were then centrifuged. The supernatant was transferred and mixed with 1mL of 100% ethanol for DNA precipitation. Samples were incubated at -20°C for 1 hour before being washed with 70% ethanol. After ethanol removal, the samples were allowed to air dry for 1 hour. DNA was subsequently resuspended in water prior to PCR. A total of 50ng of genomic DNA was used for PCR amplification.

*Fgf21* deletion was confirmed by PCR using HotStart Taq Polymerase (Ozyme) and two forwards F (5′-AGTAGGGGTCAGACGTGGTG-3′), F2 (5’-TCAGACTCAGGAGTGCAGACAA-3’) and a reverse primer R (5′-TCAGACTCAGGAGTGCAGACAA-3′) (figure 1A). Amplification conditions were as follows with Touchdown PCR : 95°C for 1min; followed by 9cycles of 95°C for 15sec, 65°C for 15sec, 68°C for 30sec and 95°C for 15sec, followed by 28cycles of 60°C for 15sec, 72°C for 1min and 72°C for 5min. This reaction produced 500-bp fragments with exons 1,2 and 3 deletion (*Fgf21* deletion) and 352-bp (*Fgf21* floxed allele).

The albumin-Cre allele was detected by PCR using the following primers pairs: Mutant forward (Alb promoter) (5′-GAAGCAGAAGCTTAGGAAGATGG -3′, Wild type forward (5′-TGGCTCGTTGTCCTTTGT -3′), and a common reverse (5′-TTGGCCCCTTACCATAACTG -3′). Amplification conditions were as follows: 95°C for 1min; followed by 35 cycles of 95°C for 15sec, 60°C for 15sec, and 72°C for 30sec ; and 72°C for 5 min. This reaction produced 390-bp (mutant), 390-bp and 530bp (heterozygote), and 530-bp (wild type) fragments.

#### DNA preparation for genotyping

DNA was extracted from tail tissue. Samples were mixed with 50µL 25 mM NaOH, and 2 mM EDTA (pH 12), then incubated for 30 min at 95°C. Samples were next, mixed, and neutralized with 50µL 40µM Tris-HCL (pH 5.0). After centrifugation, 7.6µL of supernatant was used for PCR with HotStart Taq Polymerase (Ozyme) following the manufacturer’s instructions.

### Method details

*Fasting kinetic experiments in Pparα hepatocyte-deficient and Fgf21 hepatocyte-decifient mouse model* 12-week-old male mice were fed *ad libitum* or fasted for 24h (starting at ZT0), and their glycemia and ketonemia were monitored every 4 hours. Plasma samples were prepared by centrifugation at 4,000 rpm at 4°C for 15 min. Body weight was monitored at ZT0 and ZT24 (n = 6-10 mice/genotype/experimental condition).

#### Fasting experiment in Fgf21 hepatocyte-decifient mouse model

13-week-old male and female mice were fed *ad libitum* or fasted for 20h (starting at ZT0) (n = 6-10 mice/genotype/experimental condition). Body weight, glycemia and ketonemia were measured at ZT20. After 20h of fasting, mice were sacrified by cervical dislocation. Plasma samples were prepared by centrifugation at 4,000 rpm at 4°C for 15 min. Organs and tissues were carefully collected, weighed, and frozen at -80°C until subsequent analysis. Plasma FGF21 level peaked after 20h of fasting. Therefore, we chose here to fast mice for 20h.

#### Ketogenic experiment in Fgf21 hepatocyte-decifient mouse model

10-week-old male mice were fed *ad libitum* standard diet or ketogenic diet for 9 days. On day 9, mice were sacrified by cervical dislocation (n = 10 mice/genotype/experimental condition). Body weight, glycemia and ketonemia were measured. Plasma samples were prepared by centrifugation at 4,000 rpm at 4°C for 15 min. Organs and tissues were carefully collected, weighed, and frozen at -80°C until subsequent analysis.

#### Indirect calorimetry

All animals were individually housed in a cage with lights on from 8 a.m. to 8 p.m. and an ambient temperature of 22°C ± 0.5. Mice were acclimated to their calorimetric cages for 48 h before experimental measurements. After baseline recordings for 3 days, mice were fasted for 24 h and then re-fed *ad libitum*. Data were collected every 15min. Total energy expenditure (kcal/h), oxygen consumption and carbon dioxide production (VO_2_ and VCO_2_, where V is the volume), respiratory exchange rate (RER = VCO_2_/VO_2_), and activity (counts/h) were measured using calorimetric cages (Labmaster, TSE Systems GmbH, Bad Homburg, Germany). Data analysis was carried out with Excel XP using extracted raw values of VO_2_ consumption, VCO_2_ production (mL/h), and energy expenditure (kcal/h). Subsequently, each value was expressed as a function of total lean tissue mass extracted from the EchoMRI analysis.

#### Bottle and food preference tests

All animals were individually housed in a cage. For the two-bottle sucrose preference assays, mice were allowed *ad libitum* access to standard diet and were acclimated to cages with 2 bottles of just water for 3 days. Mice were then given access to bottles with water and water containing 10% sucrose (Sigma S9378). The position of the two bottles was changed every two days to exclude position effects. Consumption was measured daily during 4 days.

For the food-preference test, feeding behaviour has been monitored using the BioDAQ system (Research Diets, Inc.). This automatic device allows continuous measurement of food intake for each animal. Before the experiment, calibration and quality control were done. For analyses, the minimum amount of food consumed by a mouse was set at 0.01 g. During the experiment, animals were singly housed at a constant temperature of 22.5 ± 1.0°C, in a reversed light/night cycle (Off: 10:30-22:30). Cages were enriched with a nesting dome. Mice had free access to water and were fed *ad libitum*. Before preference test, mice were fed with standard food (#A04, SAFE). After a 3-day acclimation period, food intake was recorded for 24 hours on the standard diet. Since each cage was equipped with two feeders, food intake consisted in the sum of both.

The 2 food-preference tests were carried out after a 20-hour fast, and was assessed over a 24-hour refeeding period, beginning at the onset of the dark period (10:30 am). For the first test, each mouse had free access to two feeders, containing either a low-protein diet (TD.90016, INOTIV) or a high-protein diet (TD.90018, INOTIV). At the end of the first test, mice were fed with standard diet again for 12 days. For the second test, feeders contained either the low-protein diet (TD.90016, INOTIV) or another high-protein diet (TD.98090, INOTIV). During tests, the distribution of diets in the right or the left feeder was randomized for each mouse.

#### Blood and tissue sampling

Prior to sacrifice, the submandibular vein was lanced and blood was collected into lithium heparin-coated tubes (BD Microtainer®, BD, Dutscher, Brumath, France). Then, mice were killed by cervical dislocation. Plasma was isolated by centrifugation (1500 × g, 15 min, 4°C) and stored at -80°C until biochemical analysis. Following sacrifice by cervical dislocation, the liver, subcutaneous white adipose tissue (scWAT), epididymal white adipose tissue (epWAT) and brown adipose tissue (BAT) were removed, weighed, and prepared for histology, or snap frozen in liquid nitrogen and stored at -80°C until use. Immediately after dissection, the pancreas was washed twice in cold HBSS 1X (without Ca^2+^ and Mg^2+^), then kept on ice during transfer to the cell culture facility. The pancreas was transferred to a sterile Petri dish containing 5 mL of HBSS 1X (without Ca^2+^ and Mg^2+^) (Gibco, 14175-095) and minced into small fragments (1–3 mm³) using sterile scissors. The fragments were transferred into a 50 mL conical tube and centrifuged at 400 × g for 2 minutes at 4°C. The supernatant was carefully discarded to remove debris and blood cells. Tissue fragments were resuspended in 5 mL of freshly prepared collagenase II solution (Collagenase II (Sigma, C6885) 2000U/mL ; HBSS 1X (with Ca^2+^ and Mg^2+^) (Gibco, 24020-117) ; HEPES (Gibco, 15630-080) 10mM ; Trypsin inhibitor (Sigma, Soybean 65035-M) 0.25mg/mL) and incubated at 37°C for 20–30 minutes. During incubation, the tissue was mechanically dissociated every 5 minutes by gentle pipetting, using pipettes of decreasing volume (25 mL, then 10 mL, then 5 mL). Once the tissue appeared homogeneously dissociated, enzymatic digestion was stopped by adding 10 mL of washing solution (HBSS 1X without Ca^2+^/Mg^2+^, 5% FBS, 10 mM HEPES). The suspension was centrifuged at 400 × g for 2 minutes at 4°C. The supernatant containing collagenase was carefully removed. The pellet was resuspended in 5 mL of washing solution and filtered through a 100 µm sterile cell strainer placed over a new 50 mL conical tube. The filter was rinsed with an additional 5 mL of washing solution. The filtrate was centrifuged at 400 × g for 3 minutes at 4°C, and the supernatant was discarded. The resulting pellet was resuspended in 1 mL of washing solution and transferred to a 1.5 mL microcentrifuge tube. A final centrifugation was performed at 450 × g for 5 minutes at 4°C. The supernatant was removed, and the pellet was immediately snap-frozen and stored at –80°C for subsequent RNA extraction.

#### Blood glucose and plasma analysis

Free fatty acids (FFAs) were determined from plasma samples using a COBASMIRA + biochemical analyzer (GenotToul Anexplo facility, Toulouse, France). Plasma glycerol was measured by enzymatic assay (Free Glycerol reagent, Sigma), blood glucose was measured with an Accu-Chek Guide glucometer (Roche Diagnostics). β-hydroxybutyrate was measured with Optium β-ketone test strips that carried Optium Xceed sensors (Abbott Diabetes Care). Plasma FGF21 was assayed using the rat/mouse FGF21 ELISA kit (Sigma) according to the manufacturer’s instructions. Free carnitine and acylcarnitines were measured from plasma (10 µL) spotted on filter membranes (Protein Saver 903 cards; Whatman), dried, and then treated as reported (Chace *et al*., 1997). Briefly, acylcarnitines were derivatized to their butyl esters and treated with the reagents of the NeoGram MSMS-AAAC kit (PerkinElmer). Their analysis was conducted using a Waters 2795/Quattro Micro AP liquid chromatography–tandem mass spectrometer (Waters, Milford, MA).

#### Gene expression

Total cellular RNA from liver, scWAT, epWAT, BAT and pancreas was extracted with Tri Reagent® (Molecular Research Center, Inc., Cincinnati, OH, USA). RNAs were quantified using a nanophotometer N60 (Implen). Total RNA samples (2 μg) were then reverse transcribed using the High-Capacity cDNA Reverse Transcription kit (Applied BiosystemsTM) for real-time quantitative polymerase chain reaction (qPCR) analyses. The primers used for the SYBR Green assays are presented in **Table 2**. Amplifications were performed by the Takyon^TM^ DNA polymerase (Takyon^TM^ low row SYBR^®^ mastermix dNTP blue kit) on a Stratagene Mx3005P thermocycler (Agilent Technology, Santa Clara, CA, USA). The qPCR data were normalized to the level of TATA-box binding protein (TBP) messenger RNA (mRNA), and analysed with LinRegPCR software (v2021.2) to determine mean efficiency (NO), which was calculated as follows: NO = threshold/(Eff meanCq), where Eff mean: mean PCR efficiency, and Cq: quantification cycle.

**Table 2:**
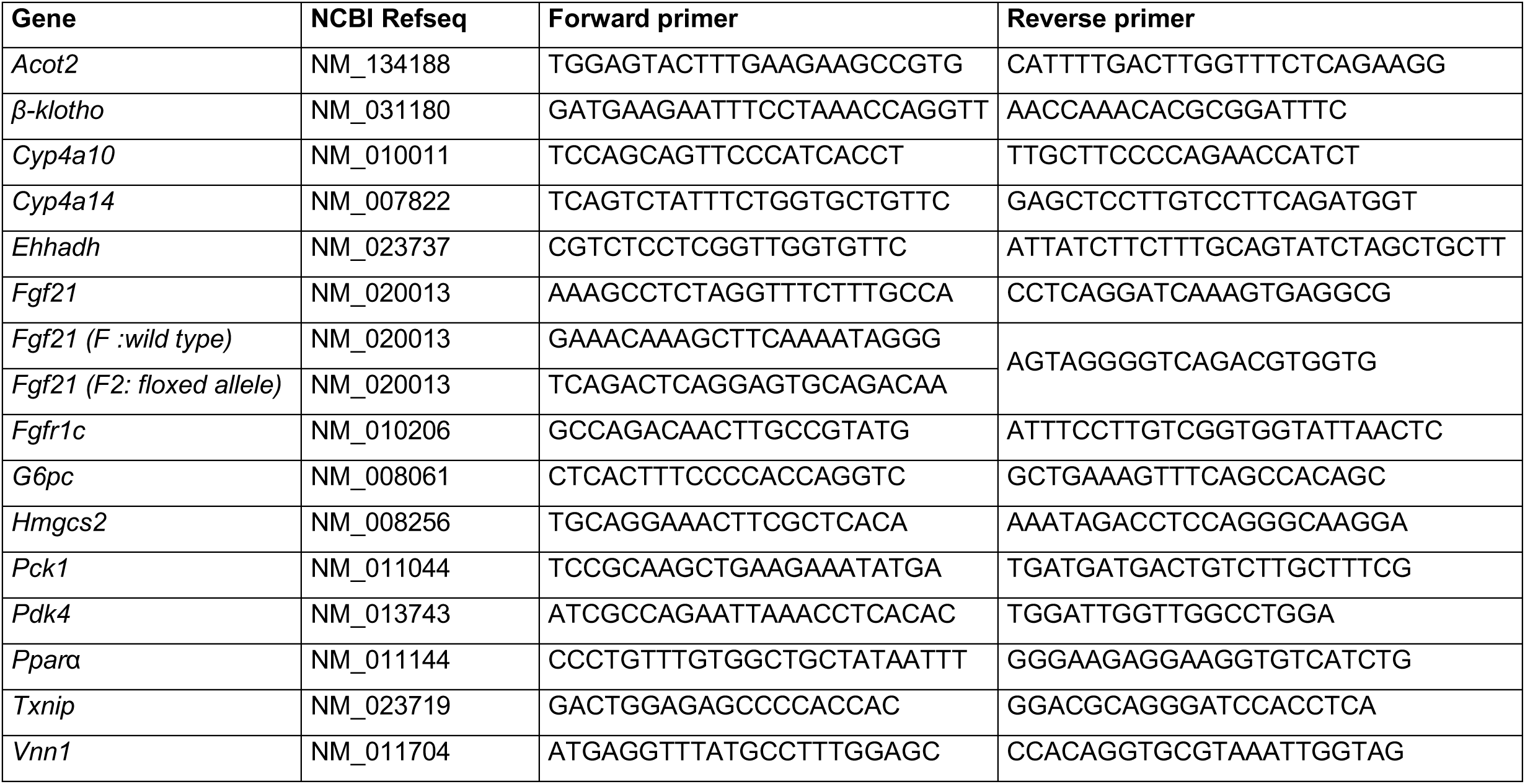
Oligonucleotide sequences for real-time qPCR.

Microarray experiments were conducted on n = 6 mice per group. Gene expression profiles were performed at the GeT-TRiX facility (GénoToul, Génopole Toulouse Midi-Pyrénées, France) using Sureprint G3 Mouse GE v2 microarrays (8 x 60K, design 074809, Agilent Technologies) following the manufacturer’s instructions. For each sample, Cyanine-3 (Cy3) labelled cRNA was prepared from 200ng of total RNA using the One-Color Quick Amp Labeling kit (Agilent technology) according to the manufacturer’s instructions. Then purification was performed by Agencourt RNAClean XP (Agencourt Bioscience Corporation, Beverly, Massachusetts, USA). Dye incorporation and cRNA yield were checked using Dropsense™ 96 UV/VIS droplet reader (Trinean, Gent, Belgium). A total of 600ng of Cy3-labelled cRNA were hybridized on the microarray slides following the manufacturer’s instructions. Immediately after washing, the slides were scanned on Agilent G2505C Microarray Scanner using Agilent Scan Control A.8.5.1 software. The fluorescence signal was extracted using Agilent Feature Extraction software v10.10.1.1 with default parameters. All experimental details and microarray data are available in NCBI’s Gene Expression Omnibus (Edgar, 2002) and are accessible through Array Express accession (E-MTAB-15848, E-MTAB-15855, E-MTAB-15880, E-MTAB-15878).

#### Histology

4% formaldehyde-fixed, paraffin-embedded liver tissue was sliced into 5 µm sections and stained with hematoxylin and eosin (H&E). The staining was visualized with a light microscope equipped with a Nikon 90i camera. 4% formaldehyde-fixed, paraffin-embedded adipose tissues were sliced into 3 µm sections and stained with hematoxylin and eosin (H&E) and scanned (GénoToul, Génopole Toulouse Midi-Pyrénées, France).

#### Liver neutral lipids analysis

Hepatic lipids were extracted as previously described (BLIGH and DYER, 1959). Briefly, tissue samples were homogenized in Lysing Matrix D tubes with 1ml methanol/5mM EGTA (ethylene glycol-bis(β-aminoethyl ether)-N,N,N’,N’-tetraacetic acid) (2:1, v/v) in a FastPrep machine (MP Biochemicals). Lipids (corresponding to an equivalent of 2 mg of tissue) were extracted in chloroform/methanol/water (2.5:2.5:2, v/v/v), in the presence of the following internal standards: glyceryl trinonadecanoate, stigmasterol, and cholesteryl heptadecanoate (Sigma-Aldrich, Saint-Quentin-Fallavier, France). Total lipids were suspended in 160 µl ethyl acetate, and the triglycerides were analyzed with gas chromatography on a Focus Thermo Electron system using a Zebron-1 Phenomenex fused-silica capillary column (5 m, 0.32 mm i.d., 0.50 µm film thickness; Phenomenex, England), as previously described (Podechard *et al*., 2018) (GénoToul Metatoul-Lipidomiqie, Génopole Toulouse Midi-Pyrénées, France). The oven temperature was programmed to increase from 200° to 350°C at a rate of 5°C/min, and the carrier gas was hydrogen (0.5 bar). The injector and the 2 detector were set to 315°C and 345°C, respectively.

#### Proton nuclear magnetic resonance (1H-NMR)-based metabolomics

Liver polar extracts were prepared and analysed using 1H-NMR-based metabolomics. All spectra were obtained on a Bruker DRX-600-Avance NMR spectrometer (Bruker) on the AXIOM metabolomics platform (MetaToul). Details on experimental procedures, data pre-treatment and statistical analysis were described previously (Lukowicz *et al*., 2018).

#### Quantification and statistical analyses

Statistical analyses on biochemical and qPCR data were performed using GraphPad Prism for Windows (version 10.00; GraphPad Software). Two-way ANOVA was performed, followed by appropriate post-hoc tests (Sidak’s multiple comparisons test) when differences were found to be significant (p < 0.05). When only 2 groups were compared, the Student’s t-test was used; p < 0.05 was considered significant. * or # p < 0.05, ** or ## p < 0.01, *** or ### p < 0.001, **** or #### p < 0.0001. Microarray data were analyzed using R and Bioconductor packages (Huber *et al*., 2015) as described in Array Express accession (E-MTAB-15848, E-MTAB-15855, E-MTAB-15880, E-MTAB-15878). Raw data (median signal intensity) were filtered, log2 transformed, corrected for batch effects (microarray washing bath and labeling serials), and normalized using the quantile method (Bolstad *et al*., 2003). A model was fitted using the limma lmFit function (Ritchie *et al*., 2015). Pair-wise comparisons between biological conditions were applied using specific contrasts. A correction for multiple testing was applied using the Benjamini-Hochberg procedure (Benjamini and Hochberg, 1995) to control the false discovery rate (FDR). Probes with an FDR <=0.05 were considered to be differentially expressed between conditions. Hierarchical clustering was applied to the samples and the differentially expressed probes, using the 1-Pearson correlation coefficient as the distance metric and Ward’s criterion for agglomeration. The clustering results are illustrated as a heatmap of expression signals. Gene ontology and transcription factor enrichment analysis were performed using Metascape (Zhou *et al*., 2019).

## Acknowledgments

We thank all members of the EZOP staff, the GeT-Trix Genotoul facility, Metatoul-Metabohub, Anexplo and Axiom facilities for their help. JB was supported by Agence Nationale de la Recherche (ANR Hepatologic ANR-21-CE14-0079-01, ANR Imagine ANR-20-CE14-0038) and Fondation pour la Recherche Médicale (Equipe FRM EQU202303016327). This work was supported by Agence Nationale de la Recherche (ANR Hepatologic ANR-21-CE14-0079-01 to HG and ANR-21-CE14-0033-01 to AB) and Fondation pour la Recherche Médicale (Equipe FRM EQU202303016327). We would like to dedicate this work to Professor David J. Mangelsdorf, in recognition of his kind support over the years, as well as his outstanding legacy and inspiration to scientists in the field.

## Author contributions

**Bruse Justine**: Conceptualization; Formal analysis; Investigation; Methodology; Visualization; Writing – original draft; Writing – review & editing. **Marbach Clothilde**: Investigation; Writing – review & editing. **Polizzi Arnaud**: Formal analysis; Investigation; Writing – review & editing. **Landre Tatiana**: Investigation; Writing – review & editing. **Salvi Juliette**: Investigation; Writing – review & editing. **Alquier-Bacquie Valérie**: Investigation; Writing – review & editing. **Chen Shiou-Ping**: Investigation; Writing – review & editing. **Huillet Marine**: Investigation; Writing – review & editing. **Rives Clémence:** Investigation; Writing – review & editing. **Martin Céline M.P.**: Investigation; Writing – review & editing. **Perrier Prunelle**: Investigation; Writing – review & editing. **Rayah Fadila**: Conceptualization; Methodology; Writing – review & editing. **Régnier Marion**: Conceptualization; Methodology; Writing – review & editing. **Weger Stefan**: Resources; Writing – review & editing. **Naylies Claire**: Investigation; Writing – review & editing. **Lippi Yannick**: Formal analysis; Supervision; Writing – review & editing. **Sommer Caroline**: Investigation; Writing – review & editing. **Albin Mikael**: Investigation; Writing – review & editing. **Lasserre Frédéric**: Investigation; Writing – review & editing. **Levade Thierry**: Resources; Supervision; Writing – review & editing. **Schupp Michael**: Resources; Writing – review & editing. **Gamet-Payrastre Laurence**: Writing – review & editing. **Kautz Léon**: Resources; Writing – review & editing. **Loiseau Nicolas**: Supervision; Writing – review & editing. **Wahli Walter**: Resources; Writing – review & editing. **Ellero-Simatos Sandrine**: Formal analysis; Investigation; Writing – review & editing. **Cruciani-Guglielmacci Céline**: Supervision; Writing – review & editing. **Montagner Alexandra**: Conceptualization; Methodology; Writing – review & editing. **Benani Alexandre**: Supervision; Funding acquisition; Writing – review & editing. **Postic Catherine**: Conceptualization; Funding acquisition; Methodology; Supervision; Writing – review & editing. **Guillou Hervé**: Conceptualization; Funding acquisition; Methodology; Project administration; Supervision; Visualization; Writing – original draft; Writing – review & editing. **Fougerat Anne**: Conceptualization; Funding acquisition; Investigation; Methodology; Project administration; Supervision; Visualization; Writing – original draft; Writing – review & editing.

## Supplementary Material

Expanded view figures

## Expanded view figure legends

**Figure EV1. Hepatocyte-specific deletion of *Fgf21* does not affect systemic responses in fasted female mice or ketogenic diet-fed male mice.** Related to Figure 1.

**(A-D)** *Fgf21* liver floxed (*Fgf21^hep+/+^*) or *Fgf21* liver knockout (*Fgf21^hep-/-^*) female mice were fed *ad libitum* or fasted for 20 hours. **(E-H)** *Fgf21* liver floxed (*Fgf21^hep+/+^*) or *Fgf21* liver knockout (*Fgf21^hep-/-^*) mice were fed a control diet (CTRL) or a ketogenic diet (KD) *ad libitum* for 9 days. **(A,E)** FGF21 plasma level was determined by ELISA. **(B,F)** Blood glucose. **(C,G)** Plasma level of β-hydroxybutyrate. **(D,H)** Body weight. Data are presented as means±SEM from 9-10 mice/group. Significance are based on 2-way ANOVA followed by Šídák’s multiple comparisons test * p<0.05; ** p<0.01; *** p<0.001; **** p<0.0001. * Fasting or KD effect.

**Figure EV2. Hepatocyte-specific deletion of *Fgf21* does not affect hepatic responses in fasted female mice or ketogenic diet-fed male mice.** Related to Figure 2.

**(A-B-E-F)** *Fgf21* liver floxed (*Fgf21^hep+/+^*) or *Fgf21* liver knockout (*Fgf21^hep-/-^*) female mice were fed *ad libitum* or fasted for 20 hours. **(C-D-G-H)** *Fgf21* liver floxed (*Fgf21^hep+/+^*) or *Fgf21* liver knockout (*Fgf21^hep-/-^*) mice were fed a control diet (CTRL) or a ketogenic diet (KD) *ad libitum* for 9 days. **(A,C)** Relative liver weight. **(B,D)** Plasma alanine aminotransferase levels. **(E,G)** mRNA relative expression of *Cyp4a10, Cyp4a14, Ehhadh, Acot2, Hmgcs2* and **(F,H)** *Pparα, Vnn1, Fgf21, Txnip, Pdk4* in liver samples measured by qRT-PCR. Data are presented as means±SEM from 8-10 mice/group. Significance are based on 2-way ANOVA followed by Šídák’s multiple comparisons test * p<0.05; ** or ## p<0.01; *** p<0.001; **** or #### p<0.0001. * Fasting or KD effect. # Genotype effect.

**Figure EV3. Hepatocyte-specific deletion of *Fgf21* does not affect glucose production during the pyruvate tolerance test.** Related to Figure 2.

**(A-B)** Pyruvate tolerance test (PTT) **(A)** and area under the curve (AUC) **(B)** representing PTT results in *Fgf21* liver floxed (*Fgf21^hep+/+^*) or *Fgf21* liver knockout (*Fgf21^hep-/-^*) male mice. Data are presented as means±SEM from 9-10 mice/group. Significance is based on 2-way ANOVA followed by Šídák’s multiple comparisons test for **(A)** and t-test for **(B)**.

**Figure EV4. Hepatocyte-specific deletion of *Fgf21* does not affect autophagy gene expression in the liver of fed or fasted mice.** Related to Figure 2.

Microarray experiment performed with liver samples from *Fgf21* liver floxed (*Fgf21^hep+/+^*) or *Fgf21* liver knockout (*Fgf21^hep-/-^*) male mice fed *ad libitum* or fasted for 20 hours (n = 6 mice/group). mRNA relative expression of *Ulk1, Ulk2*, *Wdr45, Optn, Pink1, Atg3, Atg7, Atg10, Atg14 and Nbr1* in liver samples, derived from microarray results. Data are presented as means±SEM from 6 mice/group. * p_adj_<0.05; ** p_adj_<0.01; *** p_adj_<0.001; **** p_adj_<0.0001. * Fasting effect.

**Figure EV5. Hepatocyte-specific deletion of *Fgf21* affects the expression of only a few genes in the liver during fasting.** Related to Figure 3.

*Fgf21* liver floxed (*Fgf21^hep+/+^*) or *Fgf21* liver knockout (*Fgf21^hep-/-^*) mice were fed *ad libitum* or fasted for 20 hours. **(A)** Volcano plot representing the regulated genes in (*Fgf21^hep+/+^* Fasted vs *Fgf21^hep+/+^* Fed) vs (*Fgf21^hep-/-^* Fasted vs *Fgf21^hep-/-^* Fed) mice. Green dots correspond to genes with a Log_2_FC>1 but that are non-significant (p>0.05). The red dot represents significant genes (p<0.05) with a Log_2_FC>1. **(B)** LogFC of top genes up- and **(C)** down-regulated by fasting from *Fgf21* liver floxed (*Fgf21^hep+/+^*) and *Fgf21* liver knockout (*Fgf21^hep-/-^*) in liver samples (p<0.05). **(D)** mRNA relative expression of *Fut1, Nat8, Car3, Slc13a4, Lgals1, Nkiras2, March2, Ube2l6, Tmem109, Tmem100, Myom1, Adamts14, Pigyl, Gm12*, *Zfyve21, Sfxn5, Nudt13, Lect2,* derived from microarray results. Data are presented as means±SEM from 6 mice/group. * p_adj_<0.05; ** p_adj_<0.01; *** p_adj_<0.001; **** or #### p_adj_<0.0001. * Fasting effect. # Genotype effect. **(E)** mRNA relative expression of *β-Klotho, Fgfr1c* in liver samples. Data are presented as means±SEM from 6 mice/group (A-C) and from 8-10 mice/group (E). Significance is based on 2-way ANOVA followed by Šídák’s multiple comparisons test. * p<0.05; ** p<0.01; * Fasting effect.

**Figure EV6. Hepatocyte-specific deletion of *Fgf21* does not affect hepatokine gene expression in the liver of fed and fasted mice.** Related to Figure 3.

Microarray experiment performed with liver samples from *Fgf21* liver floxed (*Fgf21^hep+/+^*) or *Fgf21* liver knockout (*Fgf21^hep-/-^*) male mice fed *ad libitum* or fasted for 20 hours. mRNA relative expression of *Angptl4, Angptl6, FetuinA, FetuinB, Gdf15, Fgl1, Lect2, Sepp1, Serpinb1a, Ctsd, Inhbe, Igf1, Igfbp1, Igfbp2, Igfbp3, Postn, Rbp4, Fst, Enho, and Tsku* in liver samples, derived from microarray results. Data are presented as means±SEM from 6 mice/group. * p_adj_<0.05; *** p_adj_<0.001; **** p_adj_<0.0001. * Fasting effect.

**Figure EV7. Hepatocyte-specific deletion of *Fgf21* does not affect fasting-induced adipose tissue gene expression.** Related to Figure 4.

*Fgf21* liver floxed (*Fgf21^hep+/+^*) or *Fgf21* liver knockout (*Fgf21^hep-/-^*) mice were fed *ad libitum* or fasted for 20 hours. **(A-C)** mRNA relative expression of *Fgf21*, *Fgfr1c,* and *β-Klotho* in sub-cutaneous white adipose (scWAT) **(A)**, epidemial white adipose tissue (epWAT) **(B)**, and brown adipose tissue (BAT) **(C)**. Data are presented as means±SEM from 7-10 mice/group (A-C). Significance is based on 2-way ANOVA followed by Šídák’s multiple comparisons test. * p<0.05 ; ** p<0.01 ; *** p<0.001. * Fasting effect. **(D-F)** Volcano plots representing the regulated genes in (*Fgf21^hep+/+^*Fasted vs *Fgf21^hep+/+^*Fed) vs (*Fgf21^hep-/-^* Fasted vs *Fgf21^hep-/-^* Fed) mice in scWAT **(D)**, epWAT **(E)** and BAT **(F)**. Green dots correspond to genes with a Log_2_FC>1 but that are non-significant (p>0.05). **(G-I)** LogFC of top genes up- and down-regulated by fasting from *Fgf21* liver floxed (*Fgf21^hep+/+^*) and *Fgf21* liver knockout (*Fgf21^hep-/-^*) in scWAT **(G)**, epWAT **(H)** and BAT **(I)** (n= 6 mice/group) (D-I).

**Figure EV8. Hepatocyte-specific deletion of *Fgf21* does not affect fasting-induced brown adipose tissue responses.** Related to Figure 4.

*Fgf21* liver floxed (*Fgf21^hep+/+^*) or *Fgf21* liver knockout (*Fgf21^hep-/-^*) mice were fed *ad libitum* or fasted for 20 hours. **(A)** Relative BAT weight. **(B)** Microarray experiment performed with brown adipose tissue (BAT) samples. Hierarchical clustering shows the definition of two gene clusters (FC > 1.5; p_adj_ < 0.05) with representation of the mean cluster profiles in each cluster for the BAT dataset. **(C-D)** Gene Ontology (GO) analysis in heatmap cluster 1 **(C)** and cluster 2 **(D)** and their respective enrichment of transcription factors **(E)**. **(F)** Representative pictures of H&E staining of BAT sections. Scale bars, 100μm. Data are presented as means±SEM from 7-9 mice/group (A) and from 6 mice/group (B-E). Significance is based on 2-way ANOVA followed by Šídák’s multiple comparisons test. * p<0.05. * Fasting effect.

**Figure EV9. Hepatocyte-specific deletion of *Fgf21* does not affect white adipose tissue lipolysis in ketogenic diet-fed male mice and fasted female mice.** Related to Figure 4.

**(A-D)** *Fgf21* liver floxed (*Fgf21^hep+/+^*) or *Fgf21* liver knockout (*Fgf21^hep-/-^*) male mice were fed a control diet (CTRL) or a ketogenic diet (KD) *ad libitum* for 9 days. **(E-H)** *Fgf21* liver floxed (*Fgf21^hep+/+^*) or *Fgf21* liver knockout (*Fgf21^hep-/-^*) female mice were fed or fasted for 20 hours. **(A,E)** Relative sub-cutaneous white adipose tisse (scWAT) weight. **(B,F)** Relative epidemial white adipose tissue (eWAT) weight. **(C,G)** Plasma free fatty acids. **(D,H)** Plasma level of glycerol. Data are presented as means±SEM from 8-10 mice/group. Significance are based on 2-way ANOVA followed by Šídák’s multiple comparisons test * or # p<0.05; ** p<0.01; *** p<0.001; **** p<0.0001. * Fasting effect. # Genotype effect. (n.d.: not detected).

**Figure EV10. Calorimetry for *Fgf21* liver floxed (*Fgf21^hep+/+^*) or *Fgf21* liver knockout (*Fgf21^hep-/-^*) male mice during 24 hours of feeding (Basal), 24 hours of fasting (Fasting), or 24 hours of refeeding (Refeeding).** Related to Figure 5.

**(A)** Locomotor activity calculated during the light and dark phases. **(B)** Rate of VO_2_ consumption normalized per lean body mass. **(C)** Rate of VCO_2_ consumption normalized per lean body mass. **(D)** Energy expenditure normalized per lean body mass. **(E)** Respiratory exchange ratio (RER). Data are presented as means±SEM from 6 mice/group. Significance is based on 2-way ANOVA followed by Bonferroni’s multiple comparisons test. * p<0.05; ** p<0.01; *** p<0.001; **** p<0.0001. * Genotype effect.

**Figure EV11. Hepatic FGF21 is required for protein preference after fasting from the first hours of feeding.** Related to Figure 5.

**(A-B)** Food intake of singly housed mice was monitored over 24 hours after 4 h, 6 h, 12 h and 24 h in mice fed *ad libitum* with a standard diet **(A)** or refed *ad libitum* with a standard diet after 20 hours of fasting **(B)**. **(C-D)** Food intake of singly housed mice was monitored over 24 hours after 4 h, 6 h, 12 h and 24 h of refeeding of mice fed *ad libitum* with a low-protein diet (low-protein/high-carb diet: 6.5% protein, 80.4% carbohydrate) and a high-protein diet (high-protein/low-carb diet: 42.6% protein, 44.3% carbohydrate) **(C)** or *ad libitum* with a low-protein diet (low-protein/high-carb diet: 6.5% protein, 80.4% carbohydrate) and a high-protein diet (high-protein/high-carb diet: 17.8% protein, 70.4% carbohydrate) **(D)**. A nonlinear regression with an exponential plateau was fitted to the data, and 95% confidence bands were plotted. Only the regression curves are shown on the graphs, with data points displayed as mean ± SEM from 5-6 mice/group.

