## Supplementary figures and images for "Hepatocyte FGF21 is not required for fasting-induced metabolic responses but guides protein appetite after energy depletion"

### EV1

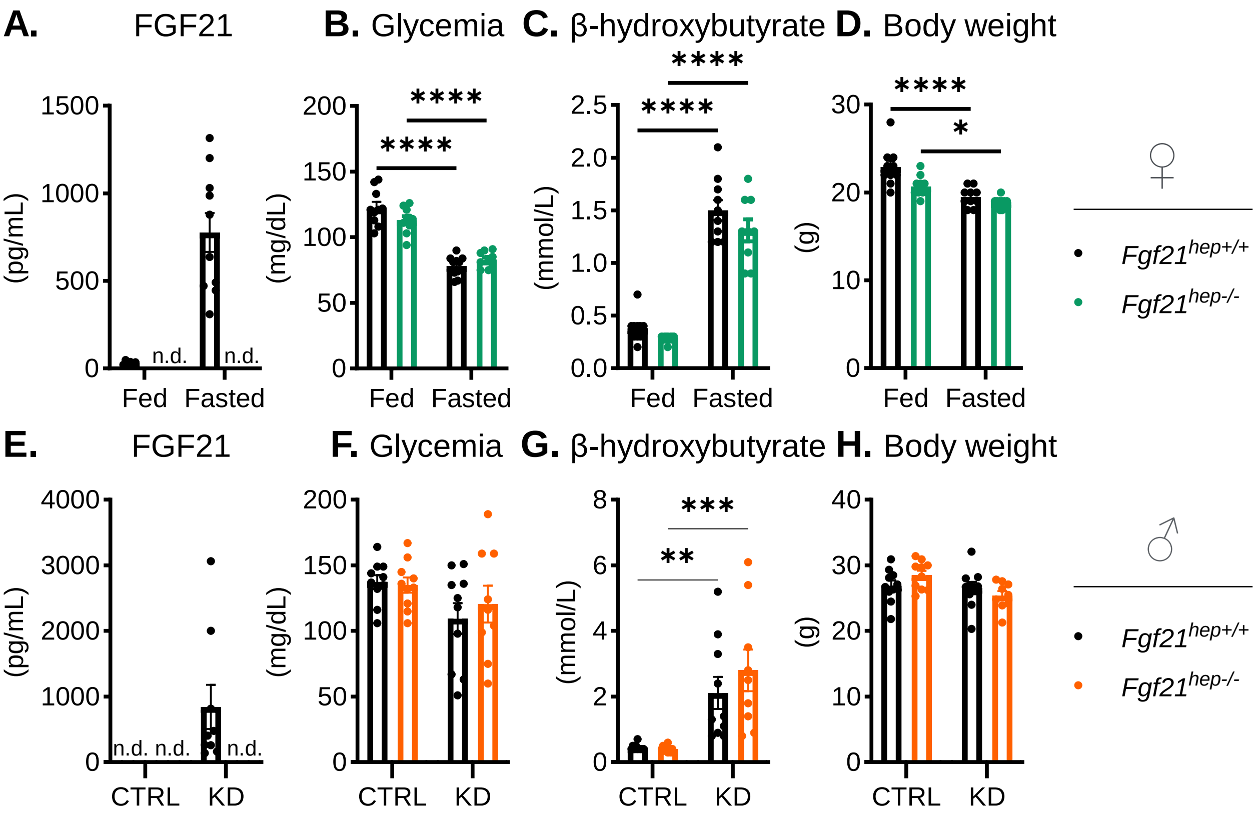

### EV2

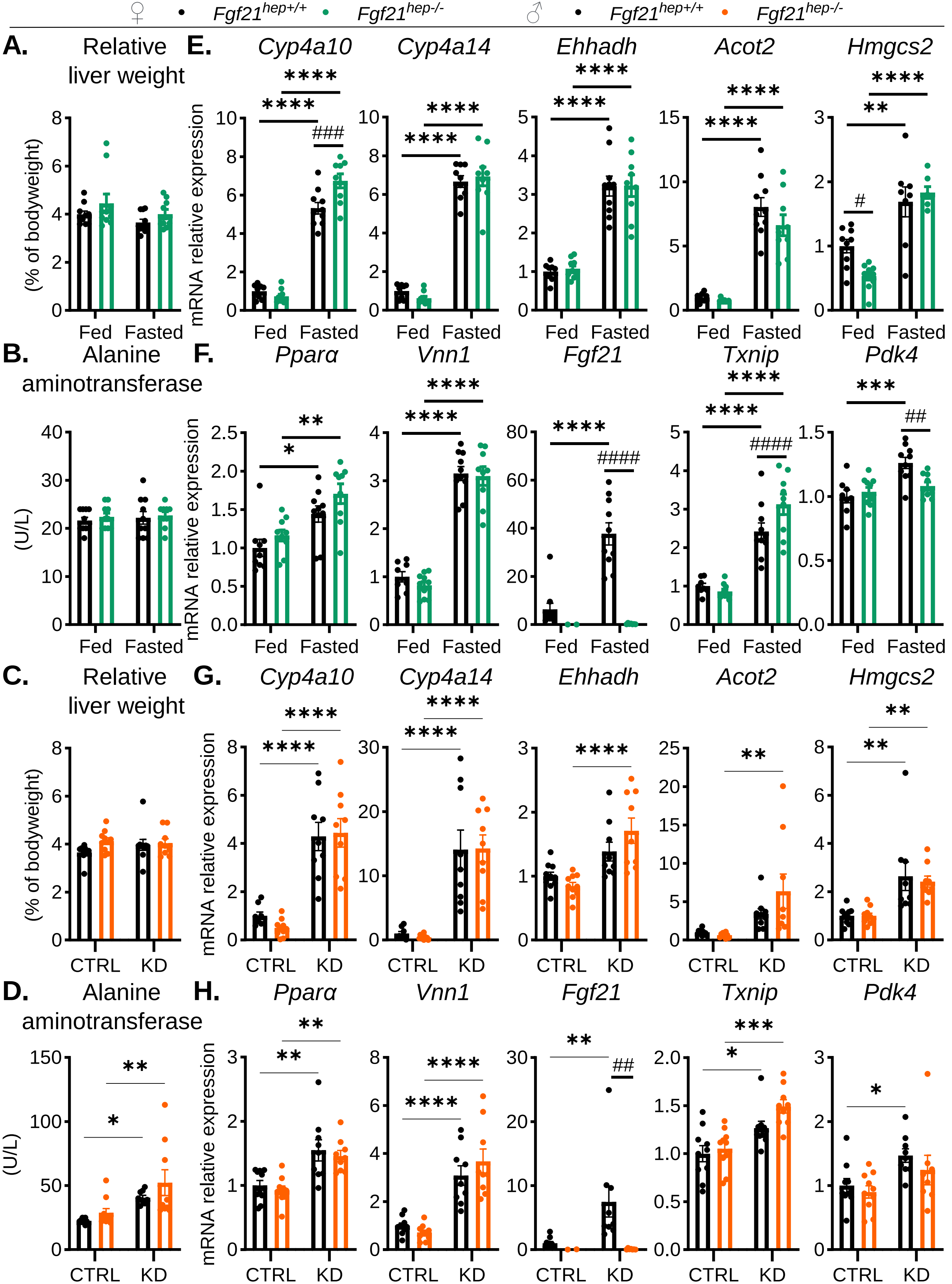

### EV3

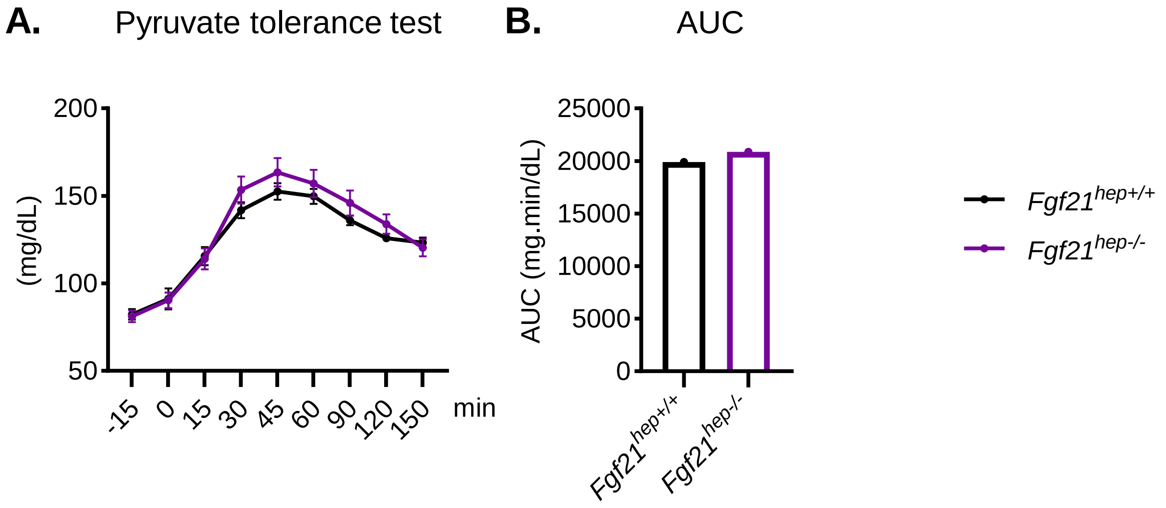

### EV4

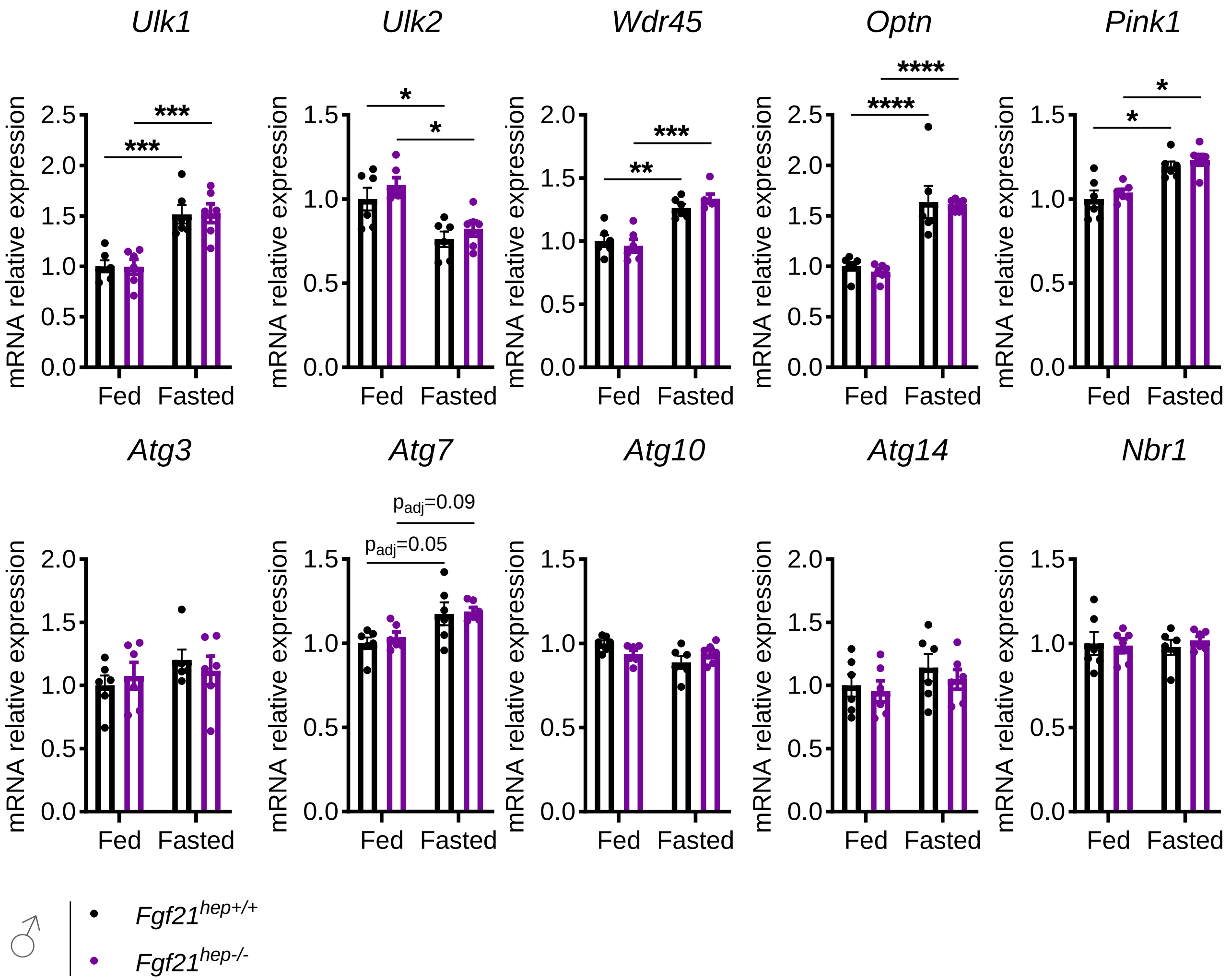

### EV5

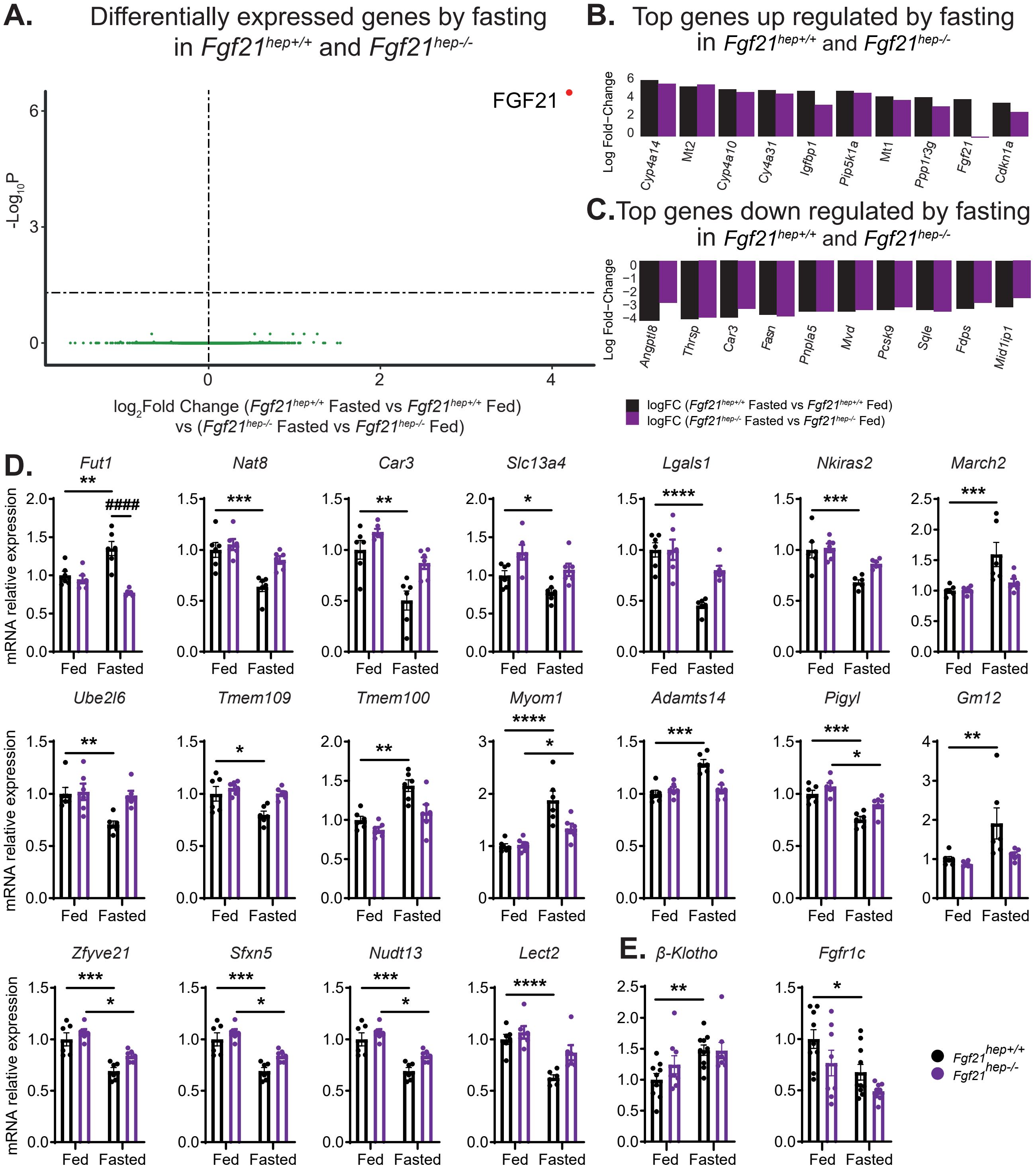

### EV6

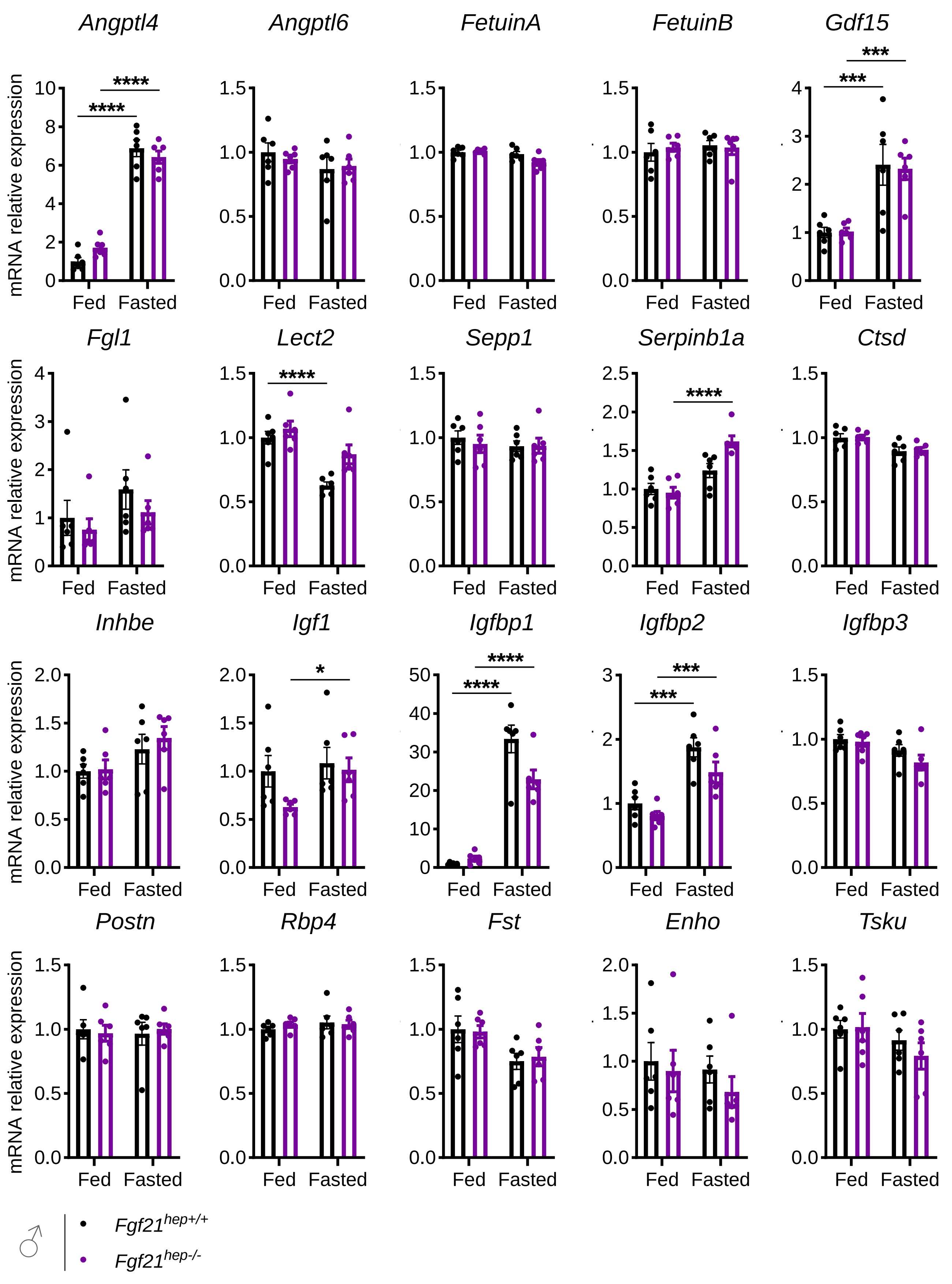

### EV7

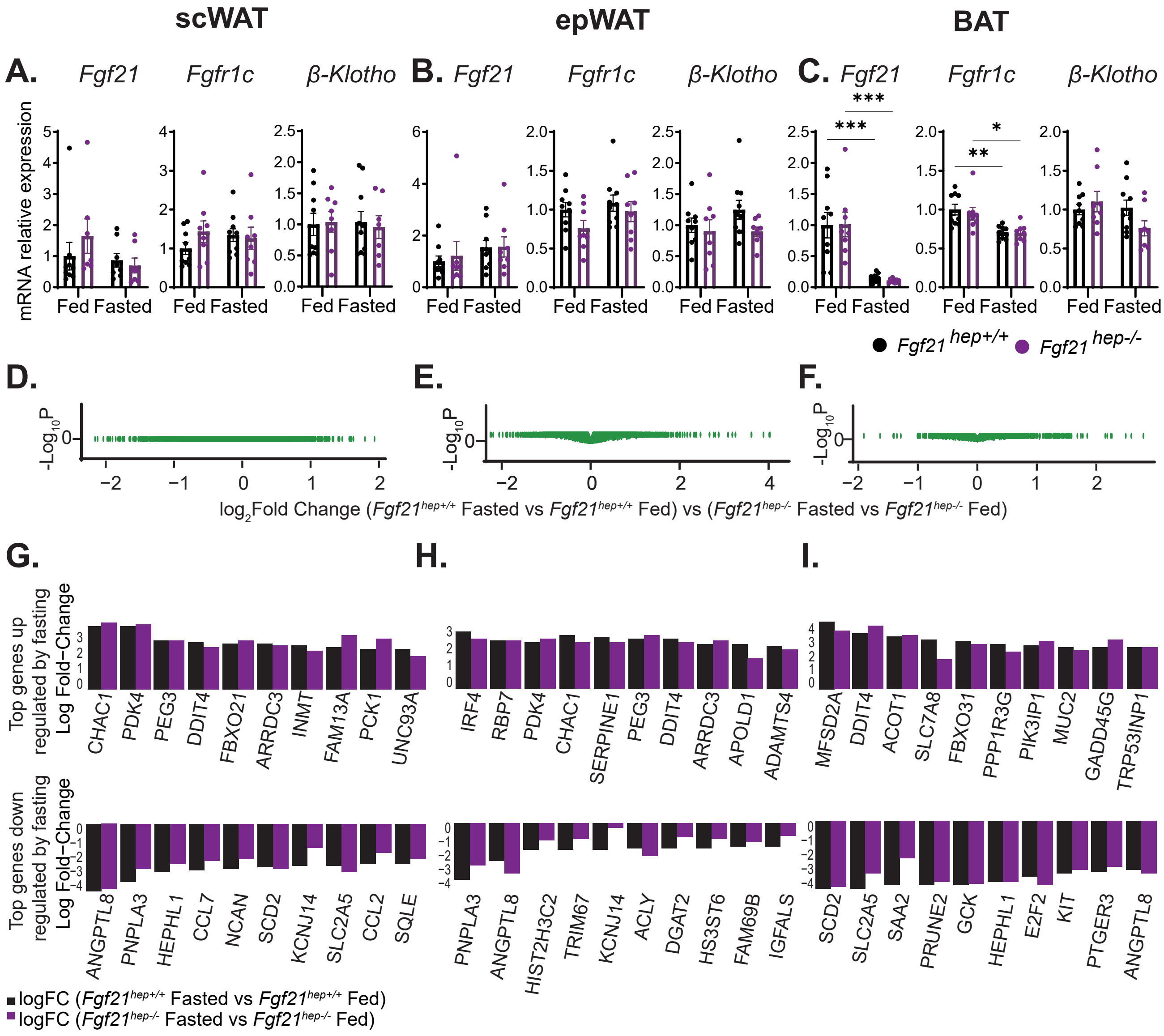

### EV8

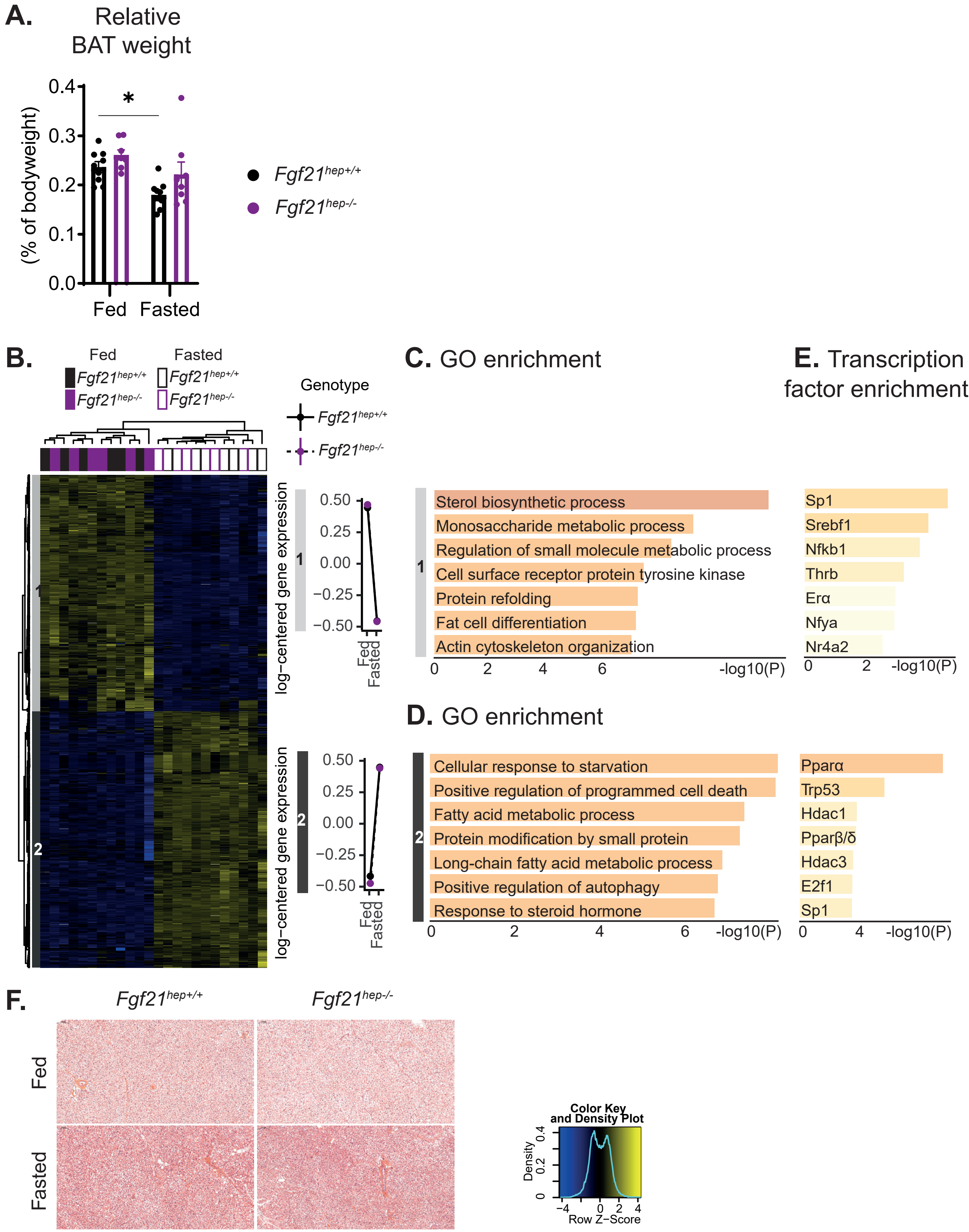

### EV9

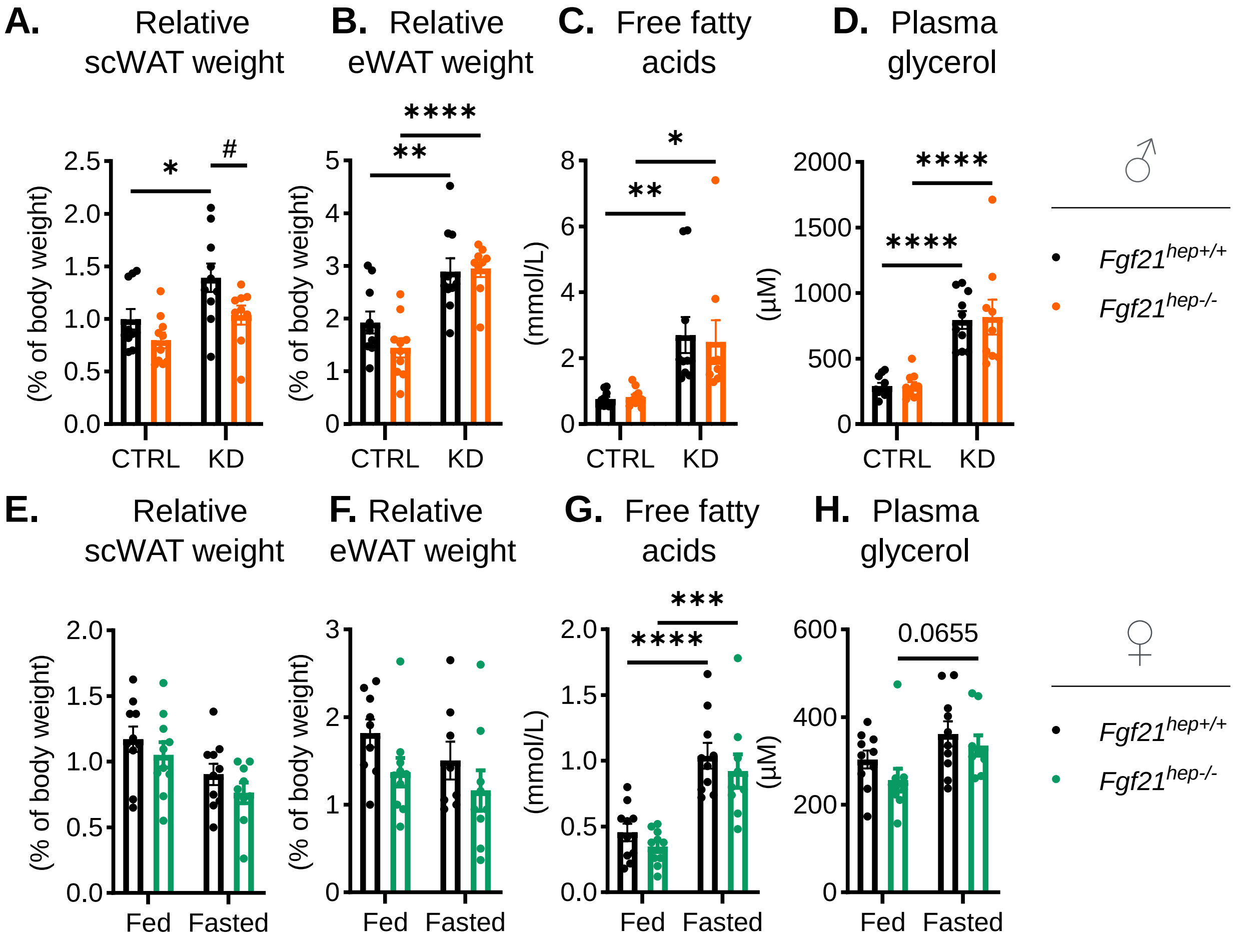

### EV10

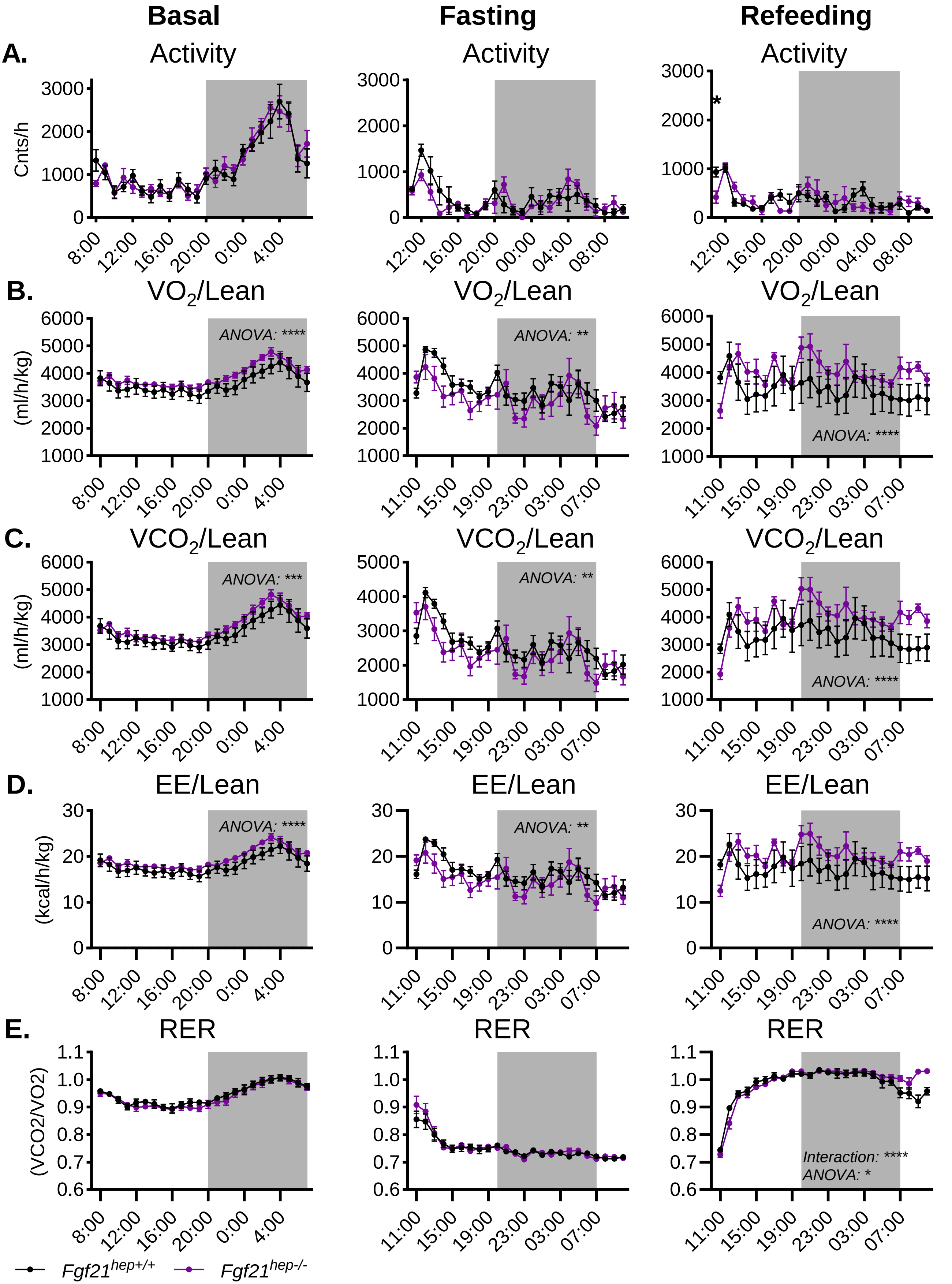

### EV11

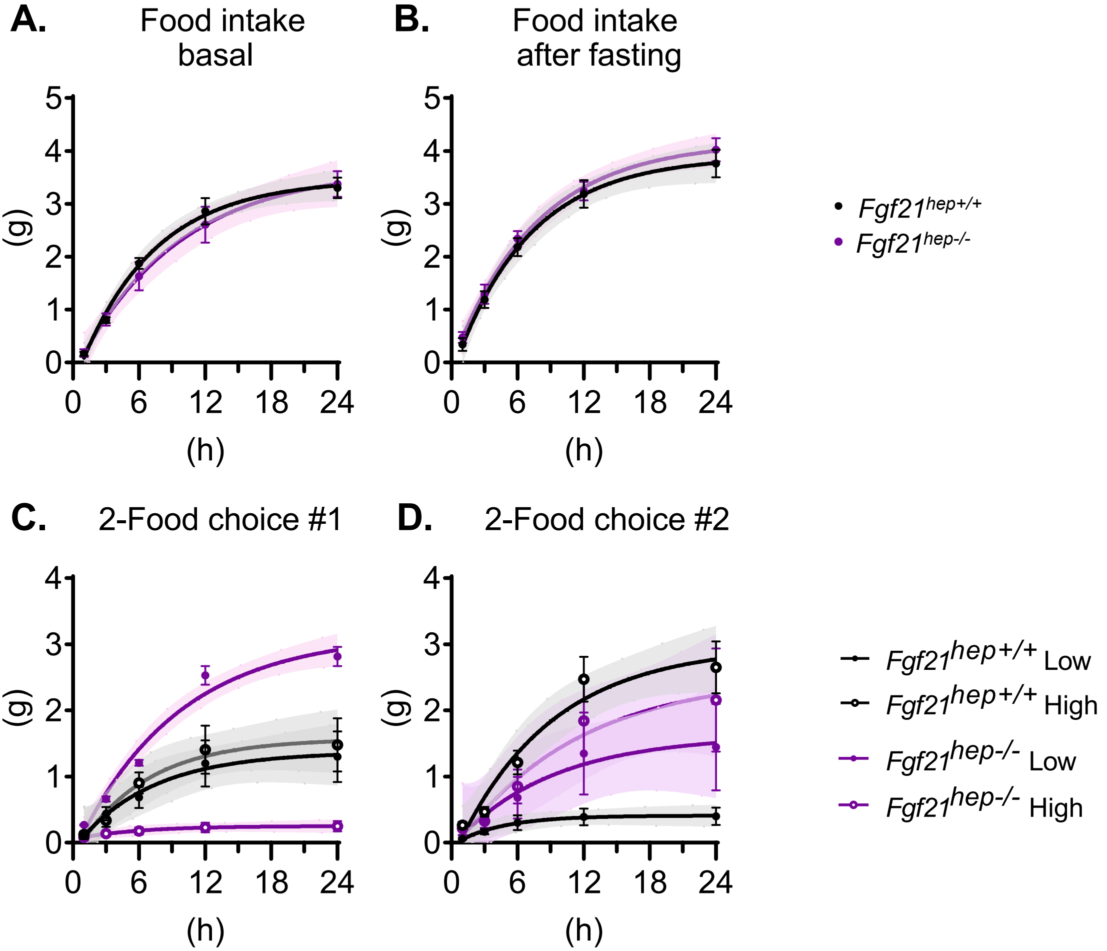
